# Biomolecular condensate viscoelasticity is dictated by the interplay between single-molecule shape memory and mesh reconfigurability

**DOI:** 10.1101/2025.10.06.680817

**Authors:** Pablo L. Garcia, Jerelle A. Joseph

## Abstract

Biomolecular condensates are membraneless organelles that compartmentalize biological functions in living cells. Formed by the phase separation of biomolecules, condensates possess a wide range of mechanical responses. However, how condensate viscoelastic responses are encoded in the chemistries of their constituents—such as intrinsically disordered proteins (IDPs)—are not well understood. Here, we employ molecular dynamics simulations to connect measurable condensate viscoelasticity to the architectural heterogeneity and dynamic reconfigurability of associative networks formed by IDPs. Using a residue-resolution coarse-grained model, we characterize biologically relevant and synthetic condensates, demonstrating that their temperature sensitivity of elasticity is sequence dependent and modeled by exponential scaling laws. We interrogate condensate mesh heterogeneity via entanglement spacing, finding that entropy-driven structural heterogeneity and reduced IDP hydrophobicity favor condensate elasticity. Furthermore, we construct graph-theoretical representations of condensates and find that interaction network topologies with an abundance of redundant node pathways translates to more load-bearing paths for mechanical stress storage. Strikingly, we discover that elastic coupling of IDPs within condensates emerges when single-molecule shape memory timescales approach meshwork reconfiguration timescales. Akin to a condensate Deborah number, this interplay of timescales for molecular and microstructural processes dictates how restoring elastic forces propagate and are stored across IDP networks; linking condensate microstructure dynamics directly to mechanical responses. Taken together, our work provides a conceptual framework of how condensates act as stress-responsive biomaterials; helping illuminate how cells exploit condensate mechanics to sense and regulate their internal environment and opening avenues for the design of condensates with programmable material properties.

**SIGNIFICANCE:** Biomolecular condensates are soft materials that exhibit diverse viscoelastic responses. We show how microstructure dynamics dictate condensate mechanics by probing the heterogeneity of local structures and how their connectivity and persistence store elastic stress. Strikingly, we discover that the ratio of timescales of single-protein shape memory to that of the underlying network rearrangement is a robust indicator of whether a condensate is dominantly viscous or elastic. This finding makes condensate mechanics predictable from sequence-dependent “Deborah numbers,” which are experimentally accessible. Our work therefore provides a tangible framework for understanding condensate stress dissipation mechanisms and opens opportunities for designing condensates with programmable material properties.

## INTRODUCTION

Multivalent macromolecules in cells actively undergo phase separation to form membrane-less organelles, also known as biomolecular condensates [1, 2]. These condensates exhibit material properties that span a continuum— from highly dynamic, liquid-like states to solid-like assemblies with arrested dynamics [3, 4]. Key properties such as fusion, wetting, viscosity (10^*−*2^ to 10^1^ Pa·s), viscoelastic stress response, and surface tension (10^*−*2^ to 10^1^ mN/m) reflect this wide rheological spectrum [5–8]. A central challenge that emerges from these observations is to elucidate how the material states of condensates are coupled to their physiological roles within the cell [9]. For instance, liquidlike condensates are readily deformable and effectively dissipate mechanical stress, suggesting a potential role in buffering mechanical perturbations. In contrast, solid-like condensates resist deformation and can store or transmit forces, potentially contributing to cellular mechanotransduction [10]. Condensates exhibiting viscoelastic behavior—characterized by a broad spectrum of relaxation times—may serve as mechanical frequency filters, selectively responding to perturbations across distinct timescales [7, 9, 11, 12].

At the molecular level, these emergent viscoelastic properties are rooted in the associative networks formed by condensate constituents [13, 14]. Among these, intrinsically disordered proteins (IDPs) with low-complexity domains (LCDs) are particularly important: their lack of stable tertiary structure and broad spectrum of conformational dynamics enable transient, multivalent interactions that assemble into mesoscale associative networks [15, 16]. Importantly, the different molecular features of IDPs may introduce varying degrees of spatiotemporal heterogeneity to condensate architectures [17–20]—which are difficult to detect and control compared to more widely studied polymer systems due to their broad dynamics [21]. For example, dynamically heterogeneous condensate architectures— such as IDP networks with regions of differing mobility, or telechelic-like structures that bridge distinct network domains [22]—are expected to shape condensate relaxation spectra, thereby modulating their ability to store, dissipate, or transmit mechanical stress [23]. In fact, IDP sequence mutations that alter condensate architectures may drive irreversible loss of condensate fluidity and transition to solidlike morphological states that are associated with pathological diseases [24, 25].

In recent years, state-of-the-art experimental methods have begun interrogating condensate mechanical responses by applying controlled perturbations and resolving the frequency-dependent phase lag between applied deformation and resulting stress. For example, combined passive and active microrheology experiments demonstrate that condensates may possess viscoelastic character akin to Maxwell fluids or Kelvin–Voigt or glassy gels [4, 7, 26]. These experiments portray how IDP sequence mutations reshape measurable condensate frequency-dependent viscoelastic spectra, change energy barriers for reconfiguration of condensate-spanning fluid networks, and alter aging dynamics [4, 27, 28]. Furthermore, atomistic force microscopy with infrared nanospectroscopy [29] and scanning probe microscopy experiments have been used to resolve how sequence modifications introduce nanomechanical heterogeneity to condensates [30, 31].

While viscoelasticity is growing in recognition as a central descriptor of condensate behavior, current advances still fall short of linking elastic properties to underlying molecular mechanisms and structure, relying instead on phenomenological interpretation. This gap limits our ability to predict or control condensate mechanics, despite their direct relevance to stability, reshaping, and mechanical signaling in cells. Consequently, recent experimental and simulation studies have begun mapping condensate microstructures [17–19, 32–34], and the interaction dynamics of their constituents [20, 35, 36] to understand how mechanical stress is stored and dissipated. Yet, there is a need to establish a framework connecting sequence-encoded interaction dynamics to emergent mesoscale architectures, and in turn, to how those architectures bear mechanical loads across different timescales.

In this work, we employ molecular dynamics (MD) simulations to connect measurable condensate viscoelastic responses to molecular-level interactions. Particularly, we present a modified version of the Mpipi coarse-grained model [37] that retains accurate phase separation propensity prediction and captures experimentally determined viscoelastic response of A1-LCD variant condensates. Beyond A1-LCD, we explore condensates formed by a diverse set of biologically relevant protein LCDs to assess how natural sequence features encode distinct condensate viscoelastic regimes. Finally, to reveal how specific physiochemical changes modulate viscoelastic response, we probe condensates formed by tyrosine-serine (Y-S) proteins, systematically varying tyrosine fraction and blockiness.

In the context of condensate life cycles (formation, dissolution and aging), we examine the temperature dependence of viscoelasticity due to protein dynamic acceleration and arrest. Our findings demonstrate that as condensates approach their critical point, their mechanical response transitions from elastic-dominated to viscous-dominated [34]. This transition is sequence-dependent and can be modeled via exponential scaling laws. Then, to elucidate the molecular origins of condensate viscoelasticity, we analyze the local geometrical features of condensates through characterizing IDP entanglement spacing—which describes the topology of meshes within condensates. Our results show that IDP sequence length, along with aromatic residue content and blockiness, influence condensate architectural heterogeneity. We also establish that delayed dynamical reconfigurability of IDP meshes, as well as enhanced cross-residue contact localization are strongly linked to condensate elastic stress response. These results provide transferrable insights into the aging mechanisms of biomolecular condensates.

As viscoelasticity arises from the distribution of mechanical loads across networks, we explore the information propagation capacity of IDP associative networks within condensates via graph-theoretical representations. We predict that condensate network topologies are highly modular, and gradually lose modularity with increasing temperature. Simultaneously, we indicate that network topology path redundancy translates into an abundance of load-bearing paths for ease of energy storage in condensates.

By employing laminar shear flow non-equilibrium simulations (NEMD), we find that condensates with heterogeneous meshes generally allow individual IDPs to adopt greater shape anisotropy, while homogeneous architectures prevent this. Surprisingly, we observe that most IDPs in the dense environment follow the coil-to-stretch transition behavior outlined in polymer theory work from De Gennes as early as 1974 [38]. This observation potentially suggests that the forces induced by shear flow screen the intermolecular forces felt by individual IDPs even in dense environments. In contrast, we observe that IDPs that form telechelic-type associative networks exhibit limits to their shape anisotropy via strain-hardening.

To establish a mechanistic interpretation of emergent elastic stress response, we compare single-molecule shape memory to mesh reconfigurability characteristic timescales. We posit that enhanced condensate elasticity is a result of single-molecule shape memory approaching the mesh reconfigurability characteristic timescale—considering both are greater than the timescale of the applied mechanical stress. Otherwise, if the single-molecule shape memory is comparatively short, IDPs undergo rapid conformational rearrangements, leading to the loss of restoring elastic forces, culminating in a viscous macroscopic response from frictional effects. As a result, we present this “condensate Deborah number” [39] as an indicator of how condensate microstructure dynamics translate to mechanical responses, which can be experimentally resolved via Förster resonance energy transfer [3] and passive microrheology experiments.

By demonstrating that condensate elasticity arises from the presence and persistence of mesh heterogeneity, as well as the capacity of individual IDPs to retain single-molecule shape memory, our framework illuminates the fundamental biophysics of elastic energy storage and force transmission in condensate associative networks. This mechanistic understanding not only advances the physical basis of condensate behavior but also lays the groundwork for rationally designing condensates with tailored functional resilience and responsiveness—offering potential applications in drug delivery, tissue scaffolding, adaptive biomaterials, and dynamic molecular gels.

## RESULTS

### Semi-flexible IDP model captures experimental biomolecular condensate viscoelastic behavior

Significant progress has been made in developing coarsegrained models that accurately capture the equilibrium thermodynamics of phase-separating IDPs. State-of-the-art coarse-grained models such as HPS [40–42], Mpipi [37, 43, 44] and CALVADOS [45–47], have linked IDP sequence composition to single-molecule properties like radii of gyration and their phase behavior. However, efforts to characterize the emergent structure [17–19, 34, 48] and dynamics [3, 35, 49, 50] of IDPs within condensed phases have only recently gained traction. Notably, recent coarsegrained models depart from the conventional treatment of IDPs as fully flexible polymers—an approach that inherently limits predictions to purely viscous material responses [51]. Instead, they adopt semi-flexible polymer representations capable of capturing viscoelastic behavior [52, 53]. This shift is supported by growing evidence that IDPs exhibit sequence-encoded mechanical properties: glycine-rich segments promote flexibility, while aromatic residues introduce stiffness [54, 55]. In parallel, single-molecule studies have demonstrated that IDPs possess sequence-dependent internal friction, which shapes their conformational dynamics in dilute conditions [56–58].

To more accurately capture the material states of condensates, we modify the Mpipi coarse-grained model Hamiltonian with the addition of a universal harmonic angle potential. We parametrize the angle potential by recapitulating the non-amino acid specific bending potential calculated from all-atom temperature replica exchange simulations of A1-LCD (wild type), using the CHARMM36m force field [59, 60] (Fig. S1). We selected the CHARMM36m force field for our simulations due to its ability to generate polypeptide backbone conformational ensembles for IDPs in quantitative agreement with NMR data such as J couplings, chemical shifts and hydrodynamic radius [61]. Furthermore, the Lennard-Jones parameters in the CHARMM36m force field have been optimized to model cation–*π* and *π*–*π* interactions [62], which are central to condensate cohesion. Finally, motivated by evidence that screened electrostatics critically influence condensate material properties [63], we also introduce temperature dependence to the Debye– Hückel screening lengths and dielectric constants into the Mpipi model [43] (see Methods). These modifications intend to correctly capture the degree of macromolecular compaction and relaxation of IDPs within condensates.

To validate our Hamiltonian modification, we confirmed that the model accurately reproduces experimentally measured radii of gyration for a diverse set of IDPs (Fig. S2), and that it captures the phase behavior of experimentally characterized A1-LCD variants [4, 64] (Fig. 1**a,b**). Importantly, we report that the Hamiltonian modification does not perturb key thermodynamic properties predicted by the original Mpipi model such as phase separation upper critical solution temperatures (Fig. S4) and surface tension (Fig. S37).

**FIG. 1:**
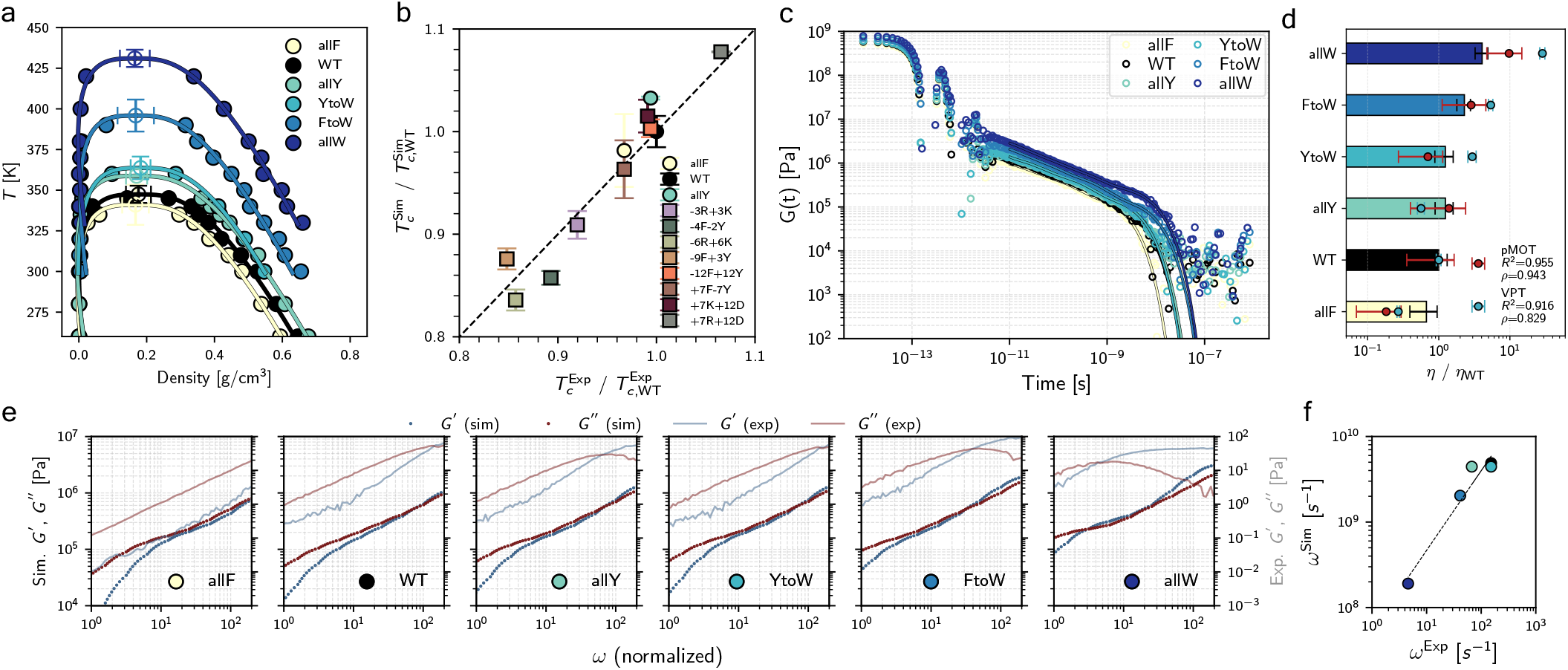
Phase separation propensity and viscoelastic behavior of A1-LCD condensates. **(a)** Phase diagrams of select A1-LCD variants with predicted critical points. Wild type A1-LCD is shown in black while mutants are shown in an increasing color gradient (F < Y < W) **(b)** Correlation between experimentally measured upper critical solution temperatures, 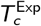 and those predicted by simulations for A1-LCD variants,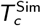 . **(c)** Shear relaxation moduli, *G* (*t*), from isotropic-box dense phase simulations at room temperature (298.15 K) show similar relaxation modes for all variants. Maxwell modes are fitted to the data to avoid noise at longer timescales. **(d)** Comparison of normalized zero-shear viscosities, *η*, from simulation to measured values from passive microrheology with optical tweezers (pMOT) and video particle tracking (VPT) experiments [4]. Pearson (*R*^2^) and Spearman (*ρ*) correlations are shown. **(e)** Loss (viscous) and storage (elastic) moduli—*G*^*′′*^ and *G*^*′*^, respectively—in comparison to experimental results over a range of normalized frequencies, *ω*. **(f)** Correlation between the experimentally measured crossover frequencies and those predicted by simulation.

While this baseline agreement with thermodynamic descriptors is essential, we seek an accurate prediction of condensate viscoelastic response. To quantify viscoelasticity in simulations, we compute the shear stress relaxation modulus *G* (*t*) through the Green–Kubo formalism with accelerated computation [65, 66]. The storage (*G*^*′*^) and loss (*G* ^*′′*^) moduli are then obtained via Schwartzian-type Fourier transforms [39] of *G* (*t*), revealing that at high frequency, *ω, G*^*′*^ dominates as IDPs are unable to relax and store deformation energy elastically, while at low *ω, G* ^*′′*^ dominates due to IDP rearrangements that dissipate energy through friction. Through direct comparisons with experiential data for A1LCD condensates [4], we find that the modified model reproduces relative zero-shear viscosities and viscoelastic spectra for these condensates (Fig.1**d–f**).

Having established that the modified Mpipi model can capture experimentally determined viscoelastic character of biomolecular condensates, we now employ it to elucidate how different IDP sequence chemistries give rise to condensates with different mechanical responses across temperature ranges.

### Temperature sensitivity of elasticity is sequence dependent

A growing body of work has established that condensate material properties are strongly influenced by the sequences of their constituent IDPs—such as aromatic residue distribution or charge blockiness [4, 36, 49, 67, 68]. Yet, despite advances in linking sequence to bulk material properties [35, 52], the microstructural basis of viscoelastic behavior remains incompletely resolved. Prior studies have shown that perturbations capable of modifying IDP networks— such as condensate desolvation [53], interaction strengthening across IDP segments to mimic inter-protein *β*-sheets formation [69], or UV-induced crosslinking [70]—can shift condensates toward more elastic, solid-like states. However, the precise ways in which condensate internal architectures are rearranged across their life cycle processes (formation, dissolution and aging) remain poorly characterized.

In this work, we conduct MD simulations controlling the magnitudes of thermal fluctuations to provide a means to induce protein dynamic acceleration and arrest. As temperature changes, certain interactions that once underpinned the condensate’s mesoscale structure lose prevalence, giving rise to new distributions of contacts that reconfigure the network into a distinct topology. This dynamic structural remodeling is expected to reflect a sequence-specific encoding of architectural plasticity, wherein an IDP’s sequence contains latent information about how the condensate reorganizes in response to thermal perturbations.

To capture the temperature-evolution of condensate viscoelasticity, we compute the loss tangent, *G* ^*′′*^*/G*^*′*^—a frequency-dependent ratio that identifies viscous (*G*^*′′*^*/G*^*′*^ *>* 1) and elastic (*G* ^*′′*^*/G*^*′*^ < 1) response regimes. For each condensate, we extract the *G* ^*′′*^*/G*^*′*^ global minima at various temperatures to characterize its maximum elastic extent (Fig. 2**a–c**). Since we are interested in examining how different condensate architectural modalities relate to viscoelastic character, we disentangle effects driven by overall interaction strength (e.g., IDPs with high aromatic content form condensates exhibiting higher upper critical solution temperatures; *T*_c_) from the effects that arise from topological remodeling by considering *G* ^*′′*^*/G*^*′*^ over a reduced temperature axis, *T/T*_c_. Through this analysis, we uncover that *G* ^*′′*^*/G*^*′*^ increases with *T/T*_c_ following exponential scaling laws (Fig. 2**c,g–i**).

**FIG. 2:**
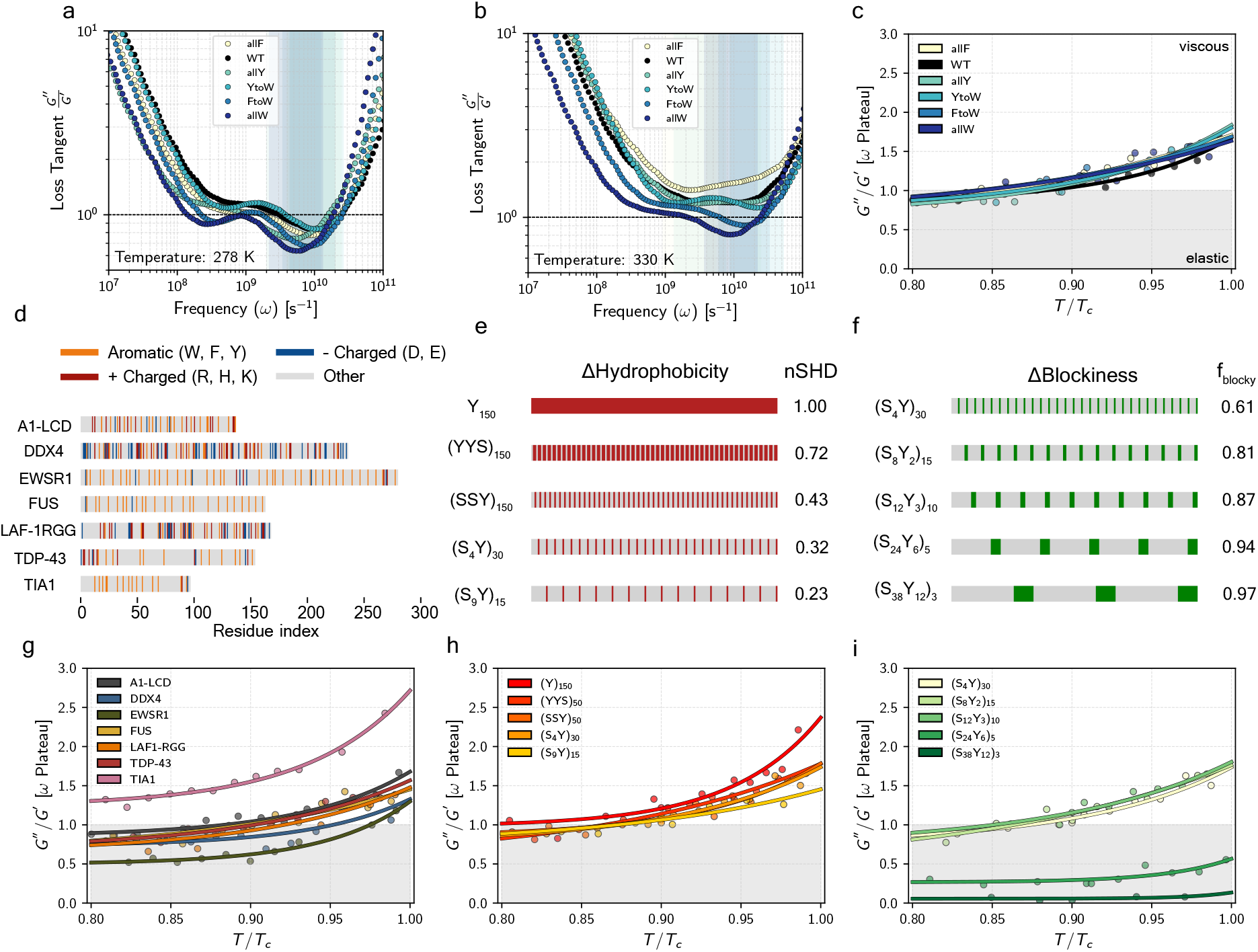
Temperature sensitivity of elasticity of biomolecular condensates. **(a, b)** Loss tangent (*G* ^*′′*^*/G*^*′*^) over a frequency range (*ω*) for two distinct temperatures. Values less than 1 represent dominance of the elastic modulus *G*^*′*^ and thus, predominant elastic character. As temperatures increases, the elastic character as well as the modes of stress relaxation (*G*^*′′*^*/G*^*′*^ local minima) are lost. Frequency “plateaus” for the local minima are highlighted. **(c)** The temperature sensitivity of elasticity of A1-LCD variant condensates. **(d)** Sequence length and composition of a set of biologically relevant LCDs. **(e)** Synthetic Y-S sequences with varying hydrophobic character captured by normalized Sequence Hydropathy Decoration (nSHD). **(f)** Synthetic Y-S sequences with varying hydrophobic blockiness at SHD ≈ 0.3. The blockiness fraction is shown. **(g–i)** The temperature sensitivity of elasticity of the set of biologically relevant LCDs, Y-S sequences with varying hydrophobic content, and Y-S sequences with different hydrophobic blockiness respectively. Color gradients display darker colors as increased hydrophobic content or increased blockiness.

Our results show that condensates formed by A1-LCD variants exhibit remarkably similar temperature sensitivity of elasticity. This suggests that, although physiochemical mutations to amino acids can substantially alter condensate viscoelastic behavior on an absolute temperature scale (Fig. 1**e**), the underlying empirical relationship between *G* ^*′′*^*/G*^*′*^ and the reduced temperature *T/T*_c_ remains relatively conserved (Fig. 2**c**). In contrast, when we examine A1-LCD alongside other biologically relevant LCDs, we observe a broader range of behaviors. Our simulations predict that condensates formed by EWSR1, DDX4, and LAF1-RGG are highly elastic, whereas those formed by TIA1 are predominantly viscous. Because these LCDs all contain fewer than ≈ 5% charged residues (Fig. 2**d**), sequence charge decoration is unlikely to be the primary factor underlying these differences. Instead, we attribute the observed trends to sequence length: longer LCD networks exhibit greater elasticity because increased chain length enhances conformational entropy, which in turn promotes entanglement effects [71]. Consistent with this picture, IDPs of roughly 150 amino acids in length (e.g., A1-LCD, FUS, TDP-43) display similar exponential scaling behaviors.

We report that our predictions from simulation are well-aligned with experimental observations. For instance, FUS condensates have been identified as Maxwell-type viscoelastic fluids through microrheology [7, 72], observed to undergo aging-associated dynamical arrest [73], and shown to posses structural and dynamical heterogeneity between kinetically trapped and fluid-like states [74]. Similarly, EWSR1 condensates exhibit markedly reduced diffusivity in the nucleolus [75] and viscoelastic behavior that can be modulated by post-translational modifications [76]. Recent computational studies also support a viscoelastic description of TDP-43’s C-terminal domain, with reported elastic response increase arising from interactions with polyA RNA [77]. While it remains challenging to capture the aging transition of TIA1 condensates into amorphous aggregates and fibrils [78], our simulations predict dynamic condensates for the wild type, consistent with FRAP analyses [79], whereas disease-linked mutations reduce mobility. However, our predictions deviate from experimental reports in the case of LAF1-RGG, as *in vitro* microrheology studies describe LAF1-RGG condensates as homogeneous and purely viscous in nature [80, 81]. These inconsistencies indicate a potential limitation in our current modeling framework or in the transferability of findings across *in vitro* platforms.

To obtain a more holistic understanding of the relationship between sequence and elasticity—and subsequently explore the exponential scaling relationships—we examine condensates formed by 150-amino-acid-long Y-S proteins, while establishing the homopolymer, Y_150_, as a baseline for comparison. We first explore the effect of varying sequence hydrophobicity, preserving the near uniform distribution of tyrosine residues noted for phase-separating prion-like domains [82], quantifying changes in hydrophobicity through the normalized Sequence Hydropathy Decoration (SHD) (see Methods). Sequence features are shown in Fig. 2**e**, with corresponding condensate elasticity extent in Fig. 2**h**. As expected, we predict higher upper critical solution temperatures for sequences with higher Y content (Fig. S7).

We observe that condensates formed by sequences with greater S content form condensates with greater elastic character and reduced sensitivity to elastic extent—noted by the decay trend in the exponential growth rate (Fig. 2**h**). We attribute this trend to reduction of energy dissipation from friction—directly captured by *G*^*′′*^: as Y residues have stronger relative cross-interaction strengths compared to S residues (Fig. S3), their diminishment in sequences leads to weaker frictional forces in the condensate environment. We then sought to capture the effect of hydrophobic patterning—or blockiness. For this analysis, we designed five sequences with equal hydrophobic content (SHD ≈ 0.3) and quantify their blockiness fraction, *f*_blocky_. Sequence features are shown in Fig. 2**f**, with corresponding condensate elasticity extent in Fig. 2**i**. We found that a small increase in blockiness did not significantly change the elasticity relative to (S_4_Y)_30_ for (S_8_Y_2_)_15_ and (S_12_Y_3_)_10_ condensates. However, (S_24_Y_6_)_5_ and (S_38_Y_12_)_3_ condensates behave like glassy solids (Fig. S12). Here, we also observe the aforementioned trend of decay in the exponential growth rate of *G* ^*′′*^*/G*^*′*^ over *T/T*_c_ space as sequence blockiness increases. We hypothesize that the decay in exponential growth rates reflects a condensate’s limited repertoire of alternative topologies upon thermal excitation, leading to a reduced capacity for architectural reorganization.

### Sequence-programmed mesh heterogeneity and reconfiguration dynamics govern condensate viscoelasticity

Elasticity in network architectures arises from the manner in which structures redistribute mechanical loads to resist deformation. Naturally, to elucidate how IDP molecular features govern condensate viscoelasticity, we must resolve their microstructures. Thus, we quantify the resultant entanglement spacing from intermolecular interactions among IDPs via correlation lengths, *ξ*. It is important to note that as per the polymer physics derivation [83], using the *ξ* order parameter assumes the system is fully isotropic and in the semi-dilute regime—prerequisites that are unsatisfied for IDP condensates. To circumvent this issue, we take inspiration from previous work that uses pore size distributions as an architectural heterogeneity metric [84], and develop a workflow that probes local entangled IDP meshes via a radius cut-off criterion (see Methods). Through this analysis, we obtain probability distribution functions, 𝒫 (*ξ*), from which we can compute coefficients of variation, *CV* (*ξ*), enabling direct comparisons of meshwork heterogeneity across condensates formed from IDPs of differing sequence composition and lengths.

As expected, we find that *CV* (*ξ*) increases with temperature from IDPs overcoming kinetic barriers and sampling wider conformational landscapes (Fig. 3**c–f**). More interestingly, different magnitudes of meshwork heterogeneity are established for different condensates. We highlight that A1-LCD variant condensates retain a relatively conserved meshwork across variants, with a very weak increase in heterogeneity with increasing aromatic residue strength (Fig. 3**c**). In contrast, our results for the set of biologically relevant LCDs demonstrate more drastic differences. For example, we determine that EWSR1 and DDX4 condensates have the most heterogeneous meshworks, while TIA1 condensates are the most homogeneous (Fig. 3**d**). Furthermore, condensates made from Y-S sequences display how decreasing sequences hydrophobicity of leads to decreased meshwork heterogeneity (Fig. 3**e**) while—in agreement with previous work [53]—we show that increasing sequence blockiness translates into increased meshwork heterogeneity (Fig. 3**f**).

**FIG. 3:**
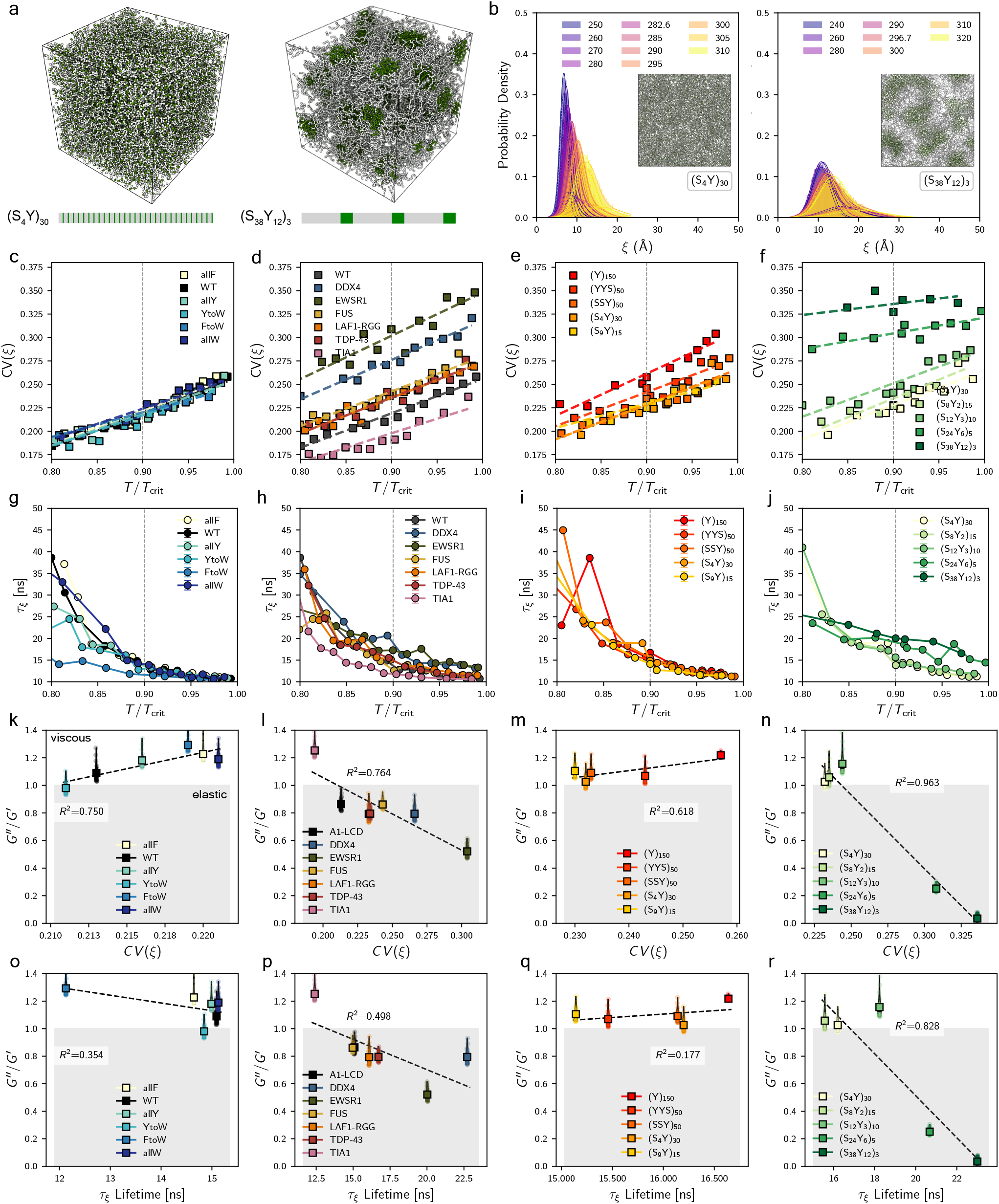
Spatial and dynamical analysis of entanglement spacing reveals connection between architectural heterogeneity and elastic stress response in condensates. **(a)** Snapshots of simulated (S_4_Y)_30_ and (S_38_Y_12_)_3_ condensates in isotropic box geometries to model dense environments. **(b)** Probability distributions of calculated entanglement spacing, *ξ*, for (S_4_Y)_30_ and (S_38_Y_12_)_3_ condensates with two fitted Gaussian functions. Insets display mesh heterogeneity. **(c–f)** Coefficient of variance, *CV* (*ξ*), for A1-LCD variants, selected biologically relevant LCDs, and Y–S sequences with varied hydrophobic content and blockiness respectively over a reduced temperature range. **(g–j)** Mesh reconfiguration lifetimes of extended mesh states, *τ*_*ξ*_. **(k–n)** Correlations between the loss tangent *G*^*′′*^*/G*^*′*^ and *CV* (*ξ*). Values of *G*^*′′*^*/G*^*′*^ less than 1 indicate dominant elastic character. **(o–r)** Correlations between the *G*^*′′*^*/G*^*′*^ and *τ*_*ξ*_.

Expecting a condensate’s deformation resistance to be tied to its architectural persistence, we investigate whether the heterogeneous meshworks are well-sustained and structurally static, or dynamic and easily reconfigurable. Towards these efforts, we calculate the characteristic timescales for mesh reconfigurability, *τ*_*ξ*_, defined as the lifetime of local entanglement spacings in the lower 25% of (*ξ*). We focus on the lower 25% as a pragmatic trade-off between selecting a clearly defined extreme tail and retaining enough events for statistically meaningful lifetime estimates; we find that results are qualitatively insensitive to modest variations of this threshold (e.g., 10–30%; Fig S28). The resulting timescale *τ*_*ξ*_ is obtained from the autocorrelation of that subpopulation and therefore quantifies how long extended neighborhoods persist before thermal fluctuations or associative rearrangements produce a topologically distinct environment. Across all our results, we show that extended local meshes are longer-lived in regimes of dynamical arrest as IDPs accept an entropic cost of narrower conformational ensembles in exchange for enthalpic gains from increased multivalency.

To establish a clear connection between how sequence-encoded IDP associative network architectures correlate with emergent condensate viscoelastic stress response, we examine the relationships between *G* ^*′′*^*/G*^*′*^ with *ξ* (Fig S27), *CV* (*ξ*) (Fig. 3**k–n**) as well as with *τ*_*ξ*_ (Fig. 3**o–r**) from simulations at 0.9*T*_c_. Our results indicate three key ideas. First, significantly decreasing IDP hydrophobicity leads to lower condensate architectural heterogeneity. We justify this observation by showing that low hydrophobicity Y-S sequences adopt collapsed conformations, which inhibits IDP crosslinking events (Fig S24). Second, more sequence blockiness leads to increased architectural heterogeneity and its longevity, which in turn, increases elastic stress response. Third, longer IDP sequences lead to more heterogeneous meshes that are also longer-lived, and strongly correlate with increased condensate elastic character.

Lastly, to understand how IDP sequence chemistry encodes accessible repertoires of network topologies, we compute intermolecular contact maps from isotropic NPT simulations at 0.9*T*_c_ (Fig. 4). Notably, condensates made from Y-S protein sequences with increasing blockiness—and consequently fewer cohesive aromatic patches—exhibit cross-contact frequencies predominantly localized to Y-rich regions. This trend is also observed for condensates formed by Y-S sequences with decreasing hydrophobicity. Hence, narrower IDP conformational ensembles arise from constraints imposed by the sustained network topologies in the condensate microenvironment. Whereas low-hydrophobicity Y-S IDPs in associative networks adopt collapsed conformations, high-blockiness IDPs experience strong entropic penalties for bridge formation imposed by local network structure. That is, IDPs become conformationally trapped upon association with two or more spatially separated large Y patches. This insight is further supported by trends in bulk mobility (Fig. S38). Previous theoretical work has termed this phenomena as chain “hopping-dominated effective diffusion” in polymer networks [85]. Thus, energy dissipation from friction is reduced from reduced IDP conformational rearrangements, leading to higher energy storage across *T/T*_c_ space (Fig. 2**i**). This is in accordance with the microscopic view of elasticity, which establishes that elasticity arises from polymeric chains resisting increasing conformational entropy. Thus, our results indicate that while absolute interaction strength primarily governs phase behavior, the sequence-encoded availability and density of strongly-interacting sequence motifs modulates the number of accessible architectures—and therefore the viscoelasticity—of condensates.

**FIG. 4:**
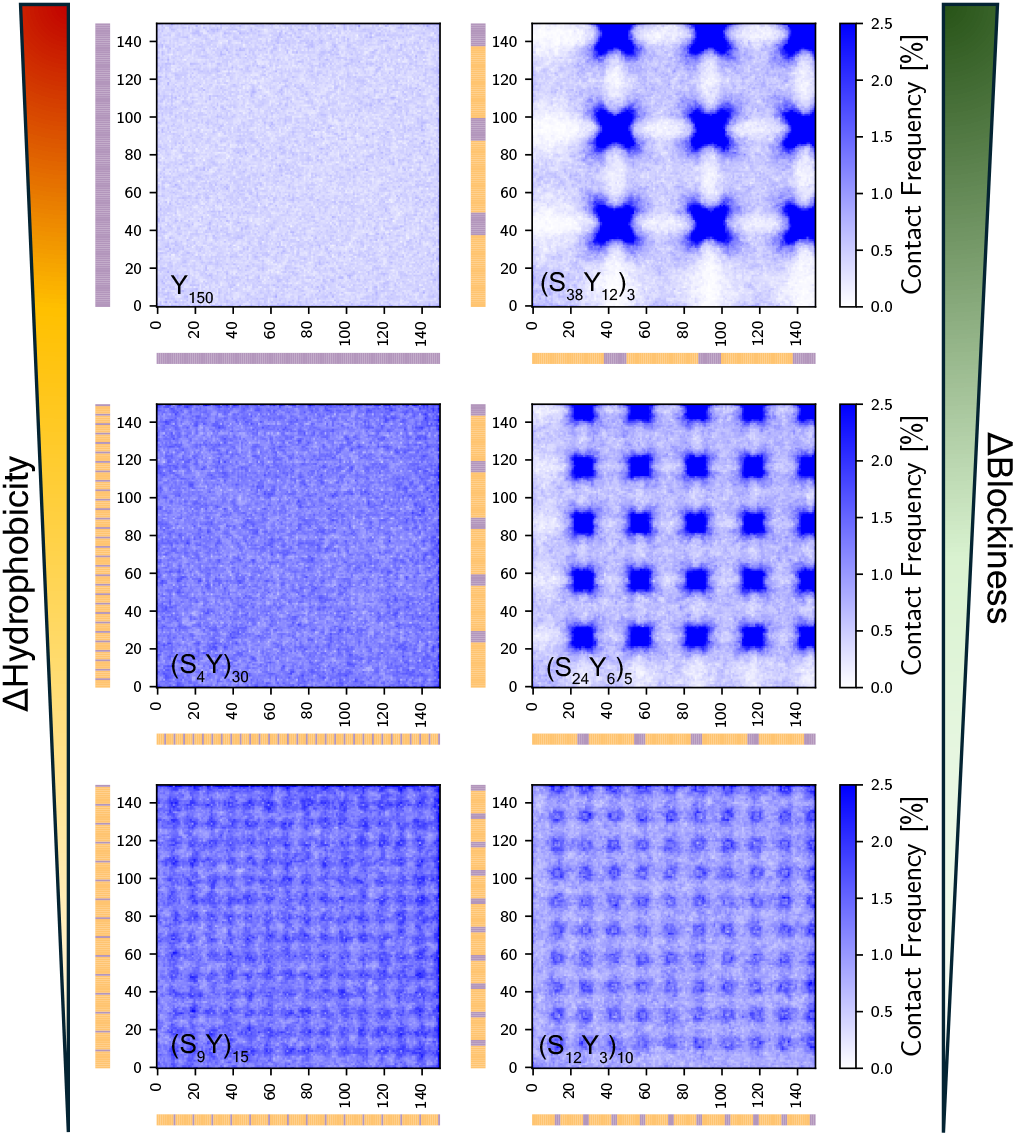
Contact maps of select Y-S variants demonstrate sequence-dependent cross-residue contact localization and conformational ensemble frustration. Y-S sequences with altered hydrophobic content show how decreasing the number of cohesive aromatic patches leads to increased cross-residue contact localization. Y-S sequences with increasing degree of blockiness display highly localized intermolecular interactions in the condensate microenvironment. This contact localization suggests narrower IDP conformational ensembles arising from network topology constraints.

### Condensate graph-network topologies reveal that node path redundancy enhances energy storage

A key aspect of characterizing viscoelastic response hinges upon identifying force transmission mechanisms within dynamic molecular assemblies like condensates. One important approach to probe the deformation-response of these IDP networks is through adopting graph-theoretical representations. Here, each IDP chain is represented as a node, and interactions across nodes as undirected edges [17]. Following our previous work [19], we construct condensate network topologies based on an a crossinteraction energy criterion of 5 *k*_B_*T* (see Methods). Among available network descriptors, the betweenness centrality, *C*_B_, of the network topologies quantifies the importance of an IDP in terms of how often it appears on the shortest paths between other IDPs interacting in distant parts of the network (Fig. 5**a, b**). This information flow description provided by *C*_B_ quantifies the number of routes for information transmission available to the network topology.

**FIG. 5:**
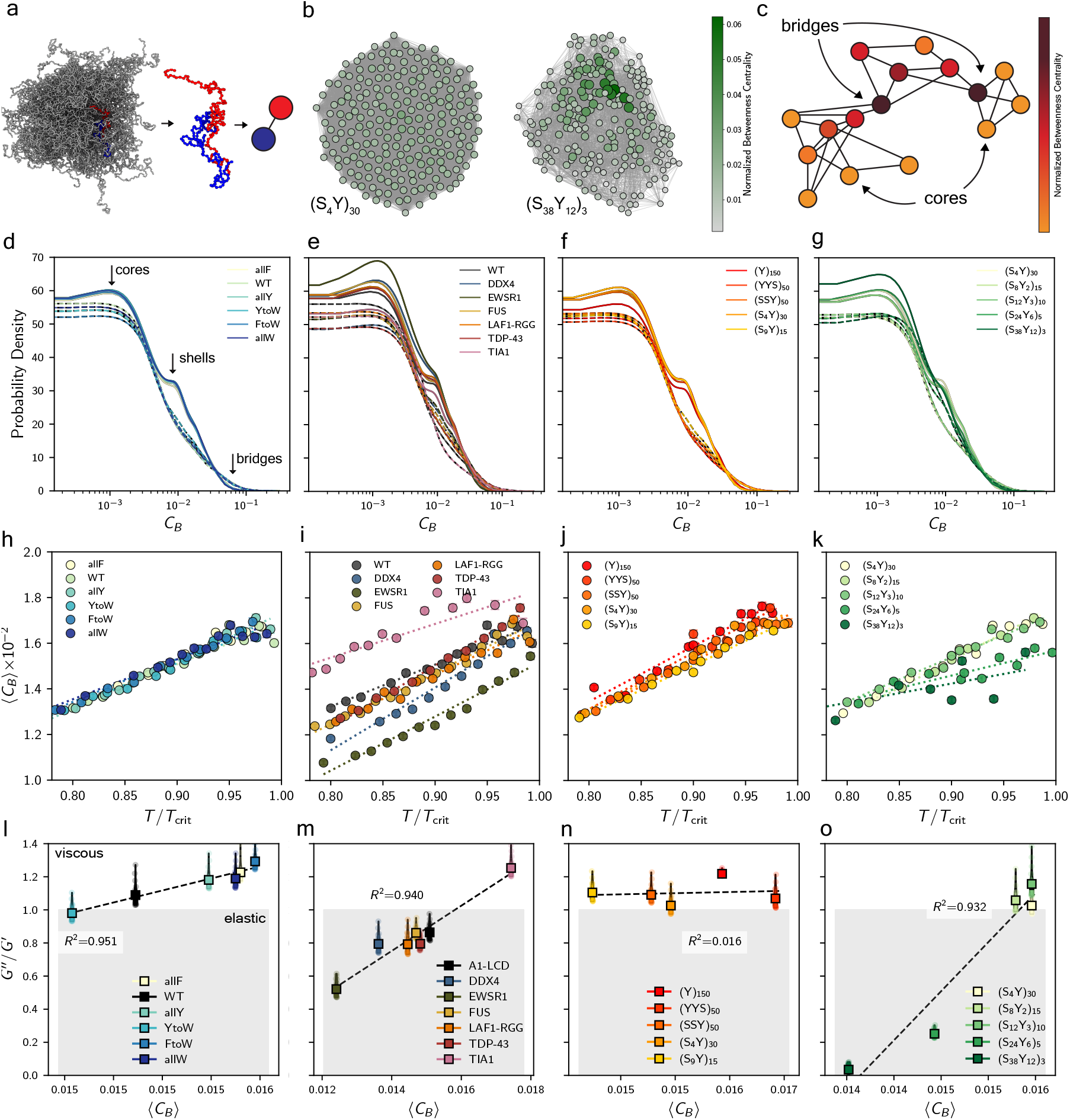
Interconnected networks that harbor an abundance of redundant load-bearing paths facilitate energy storage. **(a)** Snapshot of a biomolecular condensate explicitly showing two IDP chains in contact, which are then reduced to a two-node, one link representation. **(b)** Graph-theoretical representation of (S_4_Y)_30_ and (S_38_Y_12_)_3_ condensate dense phases respectively, produced from isotropic simulations. Nodes with higher betweenness centrality, *C*_B_, are emphasized with larger circles and darker green colors. **(c)** Cartoon representation of *C*_B_ of a modular network topology. High *C*_B_ nodes are marked by darker colors and are classified as ‘bridges’ whereas low *C*_B_ nodes are lighter and classified as ‘cores’. **(d–g)** Probability distributions of *C*_B_ at ≈ 0.8*T*_c_ (solid) and approaching criticality (0.96–0.98*T*_c_, dashed) for A1-LCD variants, selected biologically relevant LCDs, and Y–S sequences with varied hydrophobic content and blockiness, respectively. Arrows indicate different connectivity module classes. The interaction network modularity is lost as thermal fluctuation magnitudes increase. **(h–k)** Relationship between the ensemble averaged *C*_B_ and reduced temperature for A1-LCD variants, selected biologically relevant LCDs, and Y–S sequences with varied hydrophobic content and blockiness, respectively. **(l–o)** Correlations between *G*^*′′*^*/G*^*′*^ and ⟨*C*_B_⟩.

As we characterize the network topologies across different dynamical regimes, we showcase how each one moves from a role-segregated, modular structure to a more homogeneous, fast-rewiring one with increasing temperature (Figs. S29, S30, S31, S32). We present this transition through the comparison of probability distributions, 𝒫 (*C*_B_), for the highest (0.97–0.99*T*_c_) and lowest temperatures (0.8*T*_c_) in Fig. 5**d–g**. Particularly, at 0.8*T*_c_, attractions between IDPs are more selective, leading to IDP chains sorting into different connectivity module classes: ‘cores’, ‘shells’ and ‘bridges’ [86, 87]. ‘Cores’ are characterized by low *C*_B_ (they are part of many redundant paths via closed loops), ‘shells’ by intermediate *C*_B_ and ‘bridges’ by high *C*_B_ (form serial pathways for information flow).

In our previous work we demonstrated that condensate network topologies are ubiquitous for LCD-like IDPs— suggesting that the sequence design space for LCDs is vast [19]. This finding is in accordance with recent experimental evidence which suggests that LCDs do not have well-conserved sequences [88]. Here, we further provide evidence that IDP sequences that are LCD-like, despite having various inter-residue attraction modes (charged, π–π, cation–π, etc.), have identical underlying interaction network topologies described by 𝒫 (*C*_B_) (Fig. 5**d–g**). However, longer sequences such as EWSR1 lead to increased population of cores; i.e., more isolated cross-linked IDPs within their network (Fig. 5**e**).

Through the comparison between 𝒫 (*C*_B_) for the Y_150_ homopolymer and the rest of the Y-S sequences with altered hydrophobicity, we observe that modulation of sequence heterogeneity leads to changes in network topologies (Fig. 5**f**). In accordance with our contact maps (Fig. 4), the network topologies of Y-S sequences with decreased hydrophobicity possess more cores and shells—there are more local cross-links between IDPs—due to the greater contact localization to Y residues. Furthermore, we showcase a blockiness threshold in the Y-S sequences where network topologies no longer sort into three different connectivity module classes (Fig. 5**g**). Specifically, (S_38_Y_12_)_3_ and (S_24_Y_6_)_5_ condensates have underlying network topologies with one connectivity mode, as all IDPs are effectively crosslinked via the rich Y patches (Fig. S32).

To draw connections between condensate network topologies and viscoelastic stress response, we correlate the ensemble averaged betweenness centrality, ⟨*C*_B_⟩, and *G* ^*′′*^*/G*^*′*^. We know that stress relaxation requires the dissociation of cross-linked chains. Naturally, the number of cross-links per chain is expected to vary network stiffness. Indeed, our results indicate that greater redundant path count (low *C*_B_) is associated with enhanced elastic stress response for condensates(Fig. 5**l–o**). Essentially, low ⟨*C*_B_⟩ network topologies provide many alternative routes for force transmission (higher localized cross-linked IDP count) which is beneficial for energy storage. On the other hand, high ⟨*C*_B_⟩ networks possess fewer redundant node pathways, signaling diminished IDP cross-linking and formation of more short-lived contacts. These results underscore the role of IDP network connectivity as a hallmark of condensate microstructure, where harboring an abundance of redundant, loadbearing paths facilitate energy storage, ultimately enhancing the elastic stress response of condensates.

### Single-molecule shape memory and mesh reconfiguration determine condensate rheology

Classical polymer theory remarks that the stretching of polymers under flow reduces the hydrodynamic interactions between different chain segments, thus facilitating further extension in a positive feedback loop [89, 90]. This mechanism underpins a polymer’s coil-to-stretch transition, where the balance between the intramolecular forces that favor coiling and the hydrodynamic drag imposed by the flow governs the chain conformation [38]. Through this lens of a balance between disruptive and restorative forces, we sought to study how IDPs within confined condensate environments resist deformation. Here, we perform non-equilibrium molecular dynamics (NEMD) simulations at 0.9*T*_c_ to subject condensates to laminar shear flows (Fig. 6**a**).

**FIG. 6:**
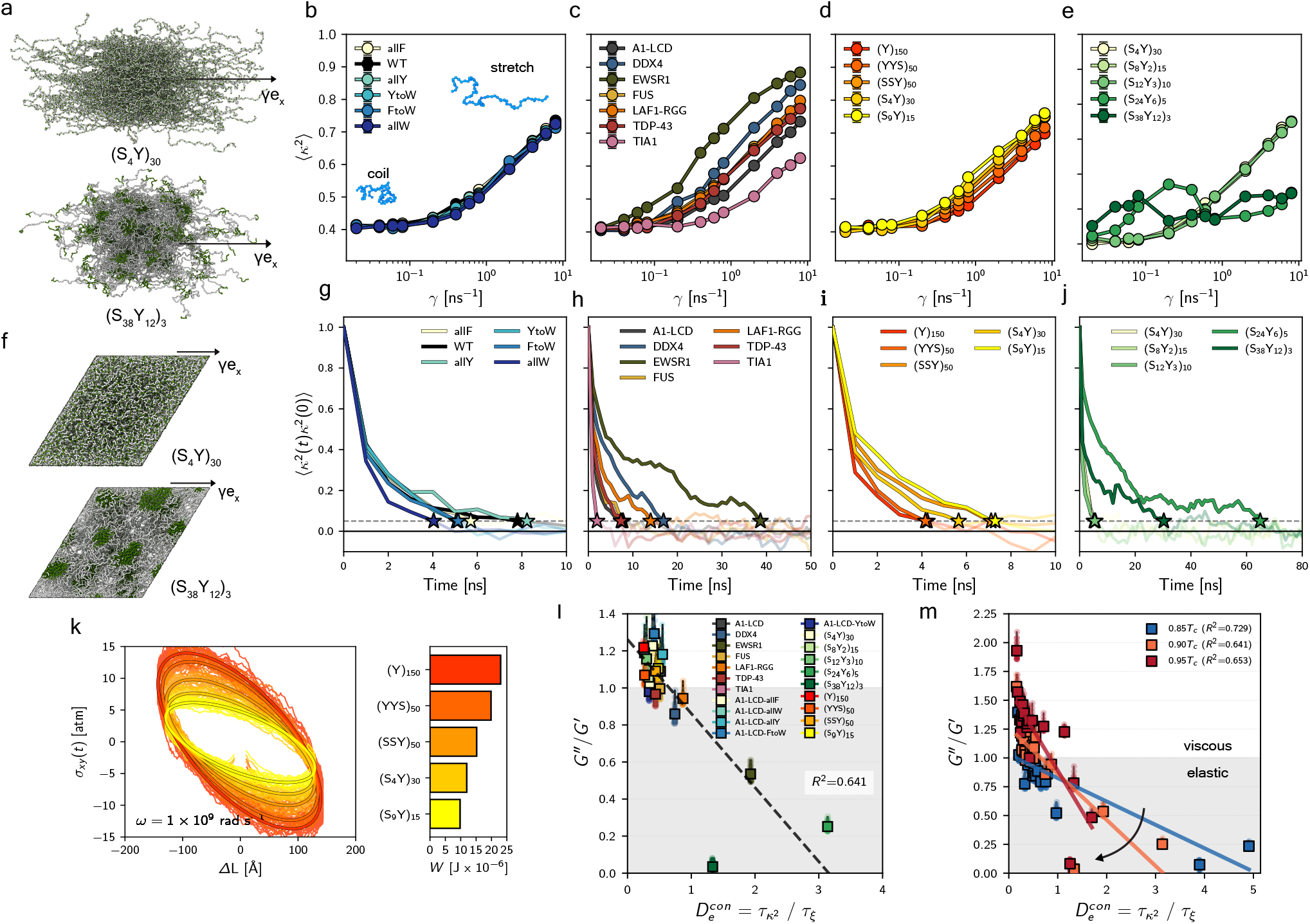
Characterization of condensate resistance to deformation reveals mechanistic insight into the origin of elastic stress response. **(a)** Simulation snapshots of unwrapped (S_4_Y)_30_ and (S_38_Y_12_)_3_ condensate dense environments undergoing constant laminar shear at 0.9*T*_c_. **(b–e)** Coil-to-stretch transitions at 0.9*T*_c_ for A1-LCD variants, the set of biologically relevant LCDs and Y–S sequences with varied hydrophobic content and blockiness respectively, marked by changes in individual IDP chain shape anisotropy, *κ*^2^. **(f)** Simulation snapshots of (S_24_Y_6_)_5_ and (S_38_Y_12_)_3_ condensates undergoing oscillatory shear in the linear viscoelastic regime at 0.9*T*_c_. **(g–j)** Autocorrelation functions of *κ*^2^ from equilibrium dense isotropic simulations for A1-LCD variants, the set of biologically relevant LCDs and Y–S sequences with varied hydrophobic content and blockiness respectively. The characteristic timescales for shape memory are shown as stars for a ⟨*κ*^2^(*t*)*κ*^2^(0) ⟩ = 0.05. **(k)** Lissajous curves and work done by the restoring elastic forces of deformed IDPs within condensates calculated from NEMD oscillatory shear simulations at 0.9*T*_c_ and a frequency of 10^*−*9^ rad s^*−*1^. Only Y–S sequences with varied hydrophobic condensates are shown to display the increase in energy dissipation from amino acid mutations that enhance cross-interaction strength. **(l)** The ratio of single-molecule shape memory over mesh reconfiguration time at 0.9*T*_c_ is related to *G*^*′′*^*/G*^*′*^ and termed here as a condensate Deborah number,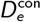. **(m)** Temperature evolution of 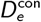. As temperatures increases, greater changes in *G*^*′′*^*/G*^*′*^ occur over condensate microstructure dynamical timescales.

To quantify the molecular-level response to shear, we track the evolution of the shape anisotropy, *κ*^2^, of individual IDPs across a range of shear rates. We calculate *κ*^2^ from the principal components of the radius of gyration tensor (see Methods). Through this characterization, we identify the extent and onset of coil-to-stretch transitions and how this transition is mediated by condensate architecture. In accordance to previous work, all characterized IDP condensates behave as shear thinning materials, whereby they exhibit lower effective viscosities compared to their zero-shear viscosities [8, 91, 92] (Fig. S33).

First, we remark that even in confined environments individual IDPs follow the coil-to-stretch transition outlined by De Gennes [38] (Fig. S34). This surprising result suggests that intermolecular interactions may be partially screened under laminar shear flow, allowing molecular conformations to be governed primarily by single-chain physics. Building on this observation, we analyze how different LCD condensates respond to shear and find that they exhibit distinct coil-to-stretch behaviors. Condensates with pronounced elastic character (e.g., EWSR1 and DDX4) permit chains to extend more readily with increasing shear rate, whereas viscous-dominated systems (e.g., TIA1 and A1-LCD) hinder chain extension (Fig. 6**c**). Moreover, condensates formed by Y–S sequences with reduced hydrophobicity facilitate chain elongation under shear (Fig. 6**d**). Finally, variants that form associative networks near the telechelic limit (e.g., (S_24_Y6)5 and (S38Y12)_3_) display strain hardening even at high shear rates, highlighting the role of network architecture in regulating single-chain dynamics (Fig. 6**e**).

Prior studies have established that the bulk mechanical relaxation times of condensates—calculated as the inverse of the crossover frequency between *G*^*′*^ and *G* ^*′′*^—are sequence dependent [26, 27, 35]. Here, we extend these observations by showing that the response of IDPs to shear is governed by the shape anisotropy decorrelation timescales, 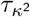, which acts as a memory kernel encoding how long past structural configurations influence present dynamics. We obtain 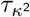 from equilibrium MD simulations, and find that condensates with larger 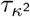 exhibit greater chain elongation under shear (Fig. 6**g–j**).

We further probe restoring elastic forces via oscillatory shear simulations at 0.9*T*_c_ (Fig. 6**f**), yielding Lissajous curves that display hallmark signatures of linear viscoelasticity and reflect stress–strain rate response [7]. In these curves, large ellipsoid aspect ratios between the box tilt, *ΔL*, and stress, *σ*_*xy*_ (*t*) correspond to increased elasticity. The energy dissipated per cycle (work, *W* ) can be calculated from area integrals, where a larger area signifies greater viscous dissipation. We first ensure that simulations are conducted in the linear viscoelastic regime (Fig. S19) by varying deformation amplitudes and frequencies. Our results for A1-LCD variant condensates demonstrate how viscous dissipation is directly related to the interaction strength of the aromatic amino acid substitution (W *>* Y *>* F), while maintaining similar viscoelasticity (Fig. S20). We also show how decreasing the hydrophobicity of Y-S sequences leads to greater elastic response and decreased viscous dissipation (Fig. 6**k**), while increasing blockiness of Y-S sequences leads to a moderate increase in elastic response with similar viscous dissipation (Fig. S20).

As shear flow drives IDPs to adopt elongated conformations, restoring elastic forces build up to favor conformational reversal. Whether these restoring forces dissipate locally or are transmitted across the IDP associative network depends on the relative timescales of molecular and microstructural processes. We propose that the ratio of 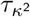 to *τ*_*ξ*_ dictates the macroscopic rheological response of condensates—akin to a Deborah number [39] for condensates: 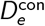. Specifically, when IDPs undergo conformational rearrangements after applied deformations more rapidly than the mesh can reorganize 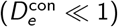, the network dissipates mechanical energy through friction. Conversely, when single-molecule memory begins to approach or exceed mesh reconfiguration timescales 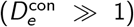, stored energy from deformation cannot be dissipated and instead, elastic coupling emerges. This enables the transmission of stored energy throughout the associative network. We present 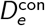, a quantifiable indicator of biomolecular condensate viscoelasticity, in Fig. 6**l** against *G* ^*′′*^*/G*^*′*^. This dimensionless quantity can be experimentally obtained by measuring single-molecule shape memory and mesh reconfiguration times through Förster resonance energy transfer [3] and passive microrheology experiments, respectively. Furthermore, we demonstrate that elastic energy storage is more sensitive to 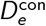 at higher temperatures (Fig. 6**m**). Together, we propose that the condensate Deborah number is a quantitative metric that directly connects microstructural dynamics of condensates to emergent mechanical response.

## DISCUSSION

Biomolecular condensates exhibit a diverse range of material states that respond differently to external forces, possibly implying distinct physiological roles inside cells. Within these dynamical assemblies, transient interactions between intrinsically disordered proteins (IDPs) give rise to percolating networks that underpin a condensate’s rheological properties. Particularly, the degree of architectural and dynamical heterogeneity of these networks may be important for distinct biological processes as they shape condensate relaxation spectra and thus modulate their ability to store, dissipate, or transmit mechanical stress [23]. Importantly, recent work suggests that single-component condensates are characterized by architecturally heterogeneous structures [17– 20, 32, 35]. Ergo, elucidating how IDP sequence features give rise to architecturally heterogeneous networks and how these relate to condensate viscoelasticity is of importance for understanding physiological phenomena involved in cellular mechanotransduction [10] and for enabling soft materials design.

In this work, we employ molecular dynamics simulations via a residue-resolution coarse-grained model to characterize the underlying viscoelastic behavior of a set of experimentally characterized A1-LCD condensates, a set of biologically relevant protein low-complexity domain (LCD) condensates (DDX4, EWSR1, FUS, LAF1-RGG, TDP-43 and TIA1), and condensates formed by binary (Y-S) sequences with altered hydrophobicity and blockiness.

We investigate how condensate viscoelasticity responds to varying thermal fluctuations, which effectively mimic different stages of the condensate life cycle by accelerating or arresting protein dynamics. Our results show that increasing thermal fluctuations reduce the ability of IDP networks to store mechanical energy, with the extent of this effect depending on sequence. This dependence follows exponential scaling laws, highlighting a quantitative framework for describing thermal sensitivity. Overall, the analysis reveals that IDPs encode latent information about how condensates reorganize under thermal perturbations. Importantly, we uncover design rules for condensate elasticity. First, from modeling A1-LCD variant condensates, we show how exchanging aromatic amino acid identities leads to negligible variations in the underlying temperature sensitivity of elasticity of condensates. Second, we observe that longer protein sequences form condensates with enhanced elasticity. Third, by analyzing Y-S protein condensates, we determine that increased sequence blockiness—such as in the case for (S_38_Y_12_)_3_, (S_24_Y_6_)_5_ or (S_9_Y)_15_—results in decreased temperature sensitivity of elasticity (Fig. 2).

Given that elastic stress response arises from how energy is transmitted across networks to resist deformation, we develop a method to characterize local geometrical features of condensate structures via the entanglement spacing order parameter, *ξ*. Through this framework, we show that the temperature evolution of condensate mesh heterogeneity (Fig. 3**c–f**) is conserved across all LCD-like condensates— condensates formed by blocky sequences are an exception. This exception arises because blocky sequences such as (S_24_Y_6_)_5_ and (S_38_Y_12_)_3_ form telechelic-like networks where Y-rich regions effectively cross-link IDPs and bridge different domains of the condensate (Fig. 4). As a result, high sequence blockiness restricts the accessible ensemble of condensate architectures across temperatures and decreases bulk mobility (Fig. S38). We also observe that condensates formed by biologically relevant LCDs with longer sequence lengths have increased mesh heterogeneity. Thus, we conclude that tuning sequence features for enhanced crosslinking capabilities—which lead to mesh heterogeneity—is a mechanism for enhancing condensate elastic response and reducing the temperature sensitivity of elasticity. Our results are in accordance with prior theoretical predictions suggesting that entropy-driven structural heterogeneity of polymeric networks is favored at the rigidity threshold of polymeric networks—or when deformation is difficult [93]. Lastly, in agreement with experiments [82], our simulation reveal that minimum sequence blockiness is crucial for maintaining fluidity in condensates.

Counterintuitively, we find that reducing sequence hydrophobicity can enhance condensate elasticity even when mesh heterogeneity diminishes (Fig. 3**e,q**). While increased heterogeneity generally correlates with stronger elastic stress storage, lowering hydrophobicity introduces a competing effect: weaker enthalpic interactions push IDPs into more compact, entropy-stabilized conformations (Fig. S24; Fig S26). These collapsed IDPs have smaller surface areas and experience lesser friction, which reduces viscous dissipation. Consequently, elastic response represents a competition between heterogeneity-driven enhancement of elastic storage and hydrophobicity-driven suppression of dissipative losses.

We probe whether mesh dynamics contribute to amplified elastic stress responses by calculating mesh reconfiguration timescales, *τ*_*ξ*_. As expected, we observe that meshes are more static in the regime of dynamical arrest, and become increasingly transient as temperature increases (Fig. 3**g– j**). Compellingly, however, notable changes in the mesh reconfiguration timescales at 0.9*T*_c_ are seen only for the biologically relevant LCDs and Y-S sequences with varying blockiness—which are the IDP subsets with most variation in mesh heterogeneity. This indicates that heterogeneous condensate meshes are longer-lived than homogeneous meshes. We then predict that condensates with longer-lived meshes have enhanced elastic behavior.

Complimenting our structural analysis, we create graph-theoretical representations of condensates to elucidate how information flow within IDP associative networks relates to energy transmission. In our previous work, we identified that the betweenness centrality, *C*_B_, of the networks represent the diverse conformational and dynamic characteristics of IDPs with greater fidelity than node degree [19]. Thus, through betweenness centrality probability distributions, 𝒫 (*C*_B_), we recognize that condensate topological networks move from a role-segregated, modular structure to a homogeneous network with increasing magnitudes of thermal fluctuations. Surprisingly, we observe that condensates whose constituents are LCD-like proteins show a similar network topology with the same connectivity module classes (Fig. 5**d–g**). Yet, IDPs with high blockiness—(S_24_Y_6_)_5_ and (S_38_Y_12_)_3_—form condensates with non-modular network topologies at any temperatures. This is representative of the aforementioned cross-linking of IDPs through the Y-rich regions that bridge the entire condensate network: all IDPs are important for condensate connectivity. Lastly, we observe that greater redundant path count (low *C*_B_) is related to increased energy storage and thus enhanced elastic response. Physically, this means that no one protein is highly important; instead, the proteins are connected with many possible routes for energy transmission.

To study how different condensates respond to mechanical stress, we perform simulations under laminar shear flow. First, in agreement with previous work on associative polymers experiencing high shear rates, our model predicts the studied condensates to be shear thinning materials [94] (Fig. S33). Then, we show that IDPs that compose viscoelastic condensates deform more readily under applied laminar shear flow (Fig. 6**b–e**). We attribute this behavior to heterogeneous meshes allowing greater IDP conformational changes. We also show how telechelic-like networks— (S_24_Y_6_)_5_ and (S_38_Y_12_)_3_—induce strain-hardening behavior against increasing laminar shear flow, indicating that the large Y patches serve as ‘anchors’ that prevent IDP deformation (Fig. 6**e**).

To understand how IDP associative networks relax and dissipate stress, we analyze the timescales of microscopic processes at different length scales. We first calculate shape anisotropy decorrelation times, 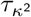, which capture singlemolecule shape memory (Fig. 6**g–j**) and quantify how long IDP dynamics remain influenced by their past conformations. We then compare 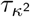 to the mesh reconfiguration time, *τ*_*ξ*_, by examining their ratio: 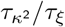. This ratio is akin to a Deborah number, *D*_e_, which is a rheological dimensionless number informing of a material’s intrinsic relaxation time against deformation process time [39]. Typically, large Deborah numbers indicate solid-like behavior. Herein, we refer to the ratio 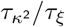 as the condensate Deborah number,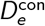.

Strikingly, 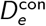 reveals how different regimes of kinetic coupling give rise to distinct mechanical responses (Fig. 6**l,m**). When individual IDPs relax faster than the mesh 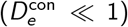, there are many opportunities for dissipative processes such as internal friction, repeated contact formation and breaking, and non-affine sliding, leading to viscous stress responses. Conversely, when conformational relaxation is governed by the lifetime of intermolecular crosslinks rather than by rapid conformational sampling 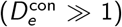 IDPs cannot relax independently but instead relax collectively, giving rise to elastic coupling across the associative network. Thus, this condensate Deborah number—which can be experimentally obtained by measuring single-molecule shape memory through Förster resonance energy transfer [3] and mesh reconfiguration times from passive microrheology—serves as a practical indicator of how condensate microstructure dynamics translate to mechanical responses.

A limitation of our work is its focus on semi-flexible disordered sequences, whereas many condensates also contain structured constituents [95]. Recent studies have shown that structured components of condensates form interaction networks stabilized by high-affinity, binding-specific interactions [96]. To approximate this effect, we use blocky Y–S IDPs, which exhibit localized, long-lived interactions. Our analysis of (S38Y12)3 and (S24Y_6_)_5_ allows us to infer that strong binding restricts accessible network architectures, imposing entropic penalties that prolong single-molecule shape memory and enhance elastic stress responses. Another limitation stems from the modified Mpipi coarse-grained model, which treats solvent implicitly and omits hydrodynamic interactions. At condensate length scales, the Reynolds number, *R*_e_ = *ρv L/µ*, is vanishingly small, indicating that longrange hydrodynamic coupling is present in this Stokes drag regime. Ignoring these interactions accelerates collective motion and underestimates viscoelastic relaxation times because solvent-driven hydrodynamic dissipation is not captured [97]. Incorporating hydrodynamics in coarse-grained models is therefore a key future direction for accurately capturing condensate dynamics and stress response.

Taken together, our work advances a mechanistic framework for condensate elasticity by (i) showing that entropy-driven mesh heterogeneity favors elastic responses, (ii) establishing network connectivity as a hallmark of load-bearing capacity, and (iii) uncovering that the balance between single-molecule shape memory and mesh reconfiguration timescales governs the extent of elastic force transmission. Building on these insights, we speculate that condensate viscoelastic responses could be dynamically tuned through post-translational modifications that reversibly expose or silence interaction motifs along IDP sequences, thereby enabling condensates to function as biosensors for stresses across defined frequency ranges. We further speculate that elastic-dominated mechanical responses may impose an additional energetic barrier that regulates molecular selectivity, rejecting molecules whose entry would require prohibitively costly mesh deformation. Together, these perspectives point to condensates as stress-responsive and compositionally selective biomaterials, with implications for understanding how cells exploit condensate mechanics to sense and regulate their internal environment and engineering condensates with programmable mechanics.

## METHODS

### Modified Mpipi model for semi-flexible protein–protein interactions

In Mpipi, each amino acid is represented by a single bead, with its corresponding physical properties such as mass, molecular diameter (*σ*), charge (*q*), and an energy scale that reflects the relative frequency of planar π–π contacts (*ε*) [37]. For the re-parametrized semi-flexible model, the total interaction energy is a sum of bonded interactions within a chain *E*_bond_, angle-interactions *E*_angle_, electrostatic contributions *E*_elec_, and short-ranged non-bonded interactions between beads *E*_pair_:

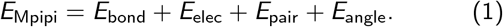

Bonds in the Mpipi model are represented via harmonic springs:

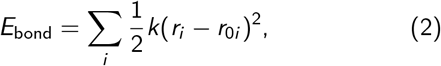

where *k* is the force constant and *r*_0i_ is the equilibrium distance. *k* is set to 0.192 kcal/mol/Å^2^ and *r*_0i_ to 3.81 Å.

In this work, the inclusion of chain stiffness is modeled through the addition of a harmonic angle potential:

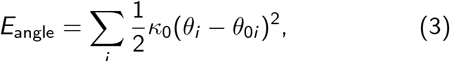

where *κ*_0_ is set to 20*k*_B_*T* and *θ*_0i_ is set to 130° to reproduce the bending potential of A1-LCD obtained from Temperature Replica Exchange atomistic simulations using the CHARMM36m [59, 60] force field in GROMACS [98].

The electrostatic contribution is modified from the original Mpipi form through a Coulomb interaction term with Debye–Hückel screening with temperature dependence:

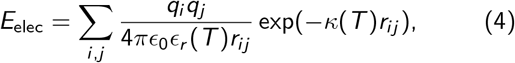

where *ϵ*_0_ is the permittivity of free space. The temperature dependence to the relative dielectric constant of water, *ϵ*_*r*_ (*T* ), follows [99]:

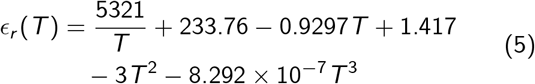

with temperature and monovalent salt concentration, *c*_*s*_ dependence to the inverse Debye–Hückel screening length, *κ* [43]:

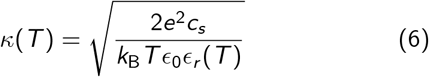

The uncharged non-bonded interactions are modeled via the Wang–Frenkel potential:

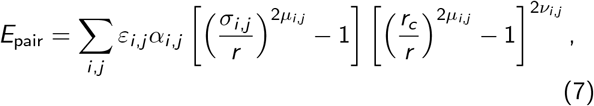

where

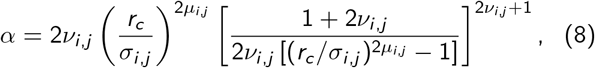

and *σ*_*i,j*_, *ε*_*i,j*_ and *µ*_*i,j*_ are parameters specified for each pair of interacting beads, and *r*_*c*_ is the cutoff radius. For most amino acids, in the original Mpipi implementation, *µ*_*i,j*_ = 1 and *ν*_*i,j*_ = 1. *r*_*c*_ is 3*σ*_*i,j*_ for each pair. All parameters for the Mpipi model can be found in Ref. [37].

### All-atom temperature replica exchange simulations

All atomistic MD simulations are conducted using the GROMACS (version May 3rd, 2024) software [98, 100]. The TIP4P water model [101] is used to solvate the proteins in cubic boxes. All Hydrogen atoms are handled by the LINCS algorithm [95] to constrain the bond lengths of hydrogen atoms to their equilibrium values. Long-range electrostatic calculations are done exclusively in GPUs via Particle Mesh Ewald [102], with a Coulomb cutoff of 1.2 nm for the solvated proteins in an ionic solution of 150 mM KCl. All protein interactions are modeled with the CHARMM36m force field [59, 60].

Initial A1-LCD structures are obtained from ColabFold v1.5.5: AlphaFold2 using MMseqs2 [103]. Subsequent minimization steps are done using the steepest descent algorithm. Then, 100 ps equilibration simulations are conducted in the canonical ensemble (*NV T* ) ensemble at 300 using 1 fs timesteps via the Verlet algorithm with Nosé– Hoover temperature coupling to preserve the natural Hamiltonian dynamics of the system [104]. Further equilibration is done in the isobaric-isothermal ensemble (*NPT* ) using the Parrinello-Rahman pressure coupling with a reference pressure of 1 atm for an additional 100 ps.

Enhanced sampling of the A1-LCD conformational ensemble is done through temperature replica exchange MD (TREMD) simulations [105]. A total of 32 replicas are simulated for 200 ns each, spanning a temperature range from 300 K to 400 K using geometric progression. Replicas are exchanged every 5 ps, achieving a recommended average exchange acceptance rate of 25–35% [106], for a cumulative simulation time of 6.4 *µ*s.

### Coarse-grained simulation of phase coexistence

All coarse-grained simulations are conducted using the LAMMPS (version 23 June, 2022) software [107–110]. Simulations rendering are performed using the Open Visualization Tool (OVITO) software [111]. The protein interactions are modeled with the Mpipi force field.

Initially, the 216 protein copies are compressed in the isothermal isobaric ensemble (*NPT* ) using a Langevin thermostat. The resultant compressed configurations are simulated in a box with 5× extended Z dimension (for a slab geometry) in the canonical ensemble (*NV T* ) at various temperatures. A simulation timestep of 10 fs is used for a total per-simulation time of 1 *µ*s.

### Determination of phase diagrams

Time-averaged density profiles can be obtained from slab simulations, where the dense and dilute phases are classified by superimposing a gaussian function on the density profile, and defining the interface location by the second derivative minimization criterion: ∂^2^*ρ/*∂*Z* ^2^ = 0.

Critical temperatures are computed using the law of coexistence densities [112]:

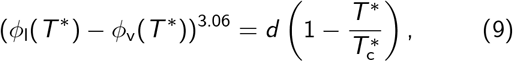

where *ϕ*_*l*_ (*T*^∗^) is the volume fraction of the high-density phase, *ϕ*_*v*_ (*T*^∗^) is the volume fraction of the low-density phase; *d* is a constant, and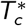 is the critical temperature. Critical densities, *ϕ*_*c*_, are computed using the law of rectilinear diameters [112]:

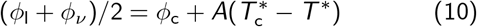

### Coarse-grained simulation of dense phase systems

Condensate dense phase simulations are set up by initially compressing 216 protein copies in the *NPT* ensemble. Subsequently, the proteins are allowed to relax to a pressure of 0 atm for 50 ns. No external pressure allows the resultant dense phase density to be solely determined by the intermolecular interactions across IDPs. Then, the proteins are equilibrated in the *NV T* ensemble for 100 ns, followed by a 1 *µ*s production run using the Nosé–Hoover thermostat [113] to preserve the natural Hamiltonian dynamics of the system [104].

### Determination of viscosity and viscoelasticity

From the MD simulations, we calculate the shear relaxation modulus, *G* (*t*), which is similar to a memory kernel of a generalized Langevin particle, and measures how shear stress relaxes upon the introduction of unit-step strain at t=0 [51, 68]. It is useful to mention that for elastic solids, *G* (*t*) approaches a constant—that is, shear stress never relaxes—whereas in viscous liquids, *G* (*t*) is a delta function of time. More importantly, in viscoelastic liquids, shear relaxation occurs at finite rates, particularly expressed through exponentially decaying functions of time, which are referred to as Maxwell modes.

In our simulations, *G* (*t*) is calculated through the Green– Kubo formalism for isotropic systems in the *NV T* ensemble using the autocorrelation function of the off-diagonal components of the pressure tensor, *σ*_*αβ*_:

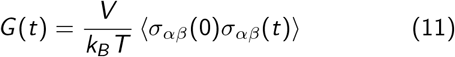

A more accurate expression for *G* (*t*) can be found directly using the six independent components of the pressure tensor [66, 114]:

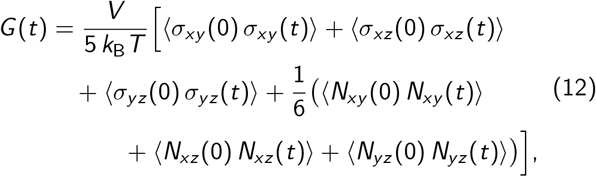

where the normal stress tensors are *N*_*αβ*_ = *σ*_*αα*_ *σ*_*ββ*_. This is done in LAMMPS using the compute `ave/correlate/long` from the USER-MISC package [115].

From here, condensate zero-shear viscosity, *η*, can be calculated as follows:

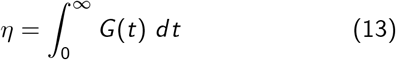

To avoid the noise at the terminal decay region, and thus accurately determine the time integral of the shear stress modulus, we separate the range of frequencies into an analytical fit at short frequencies and a series of Maxwell modes at long frequencies after some time *t*_0_ when intramolecular oscillations of *G* (*t*) have decayed:

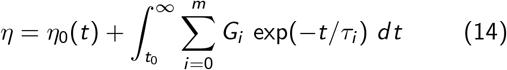

We identify the viscous and elastic moduli of the IDP condensates, by performing a Schwartzian-type Fourier transform [39] of the *G* (*t*) for each isotropic simulation of the condensates:

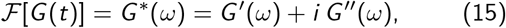

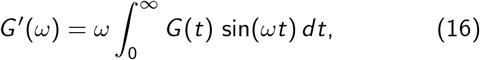

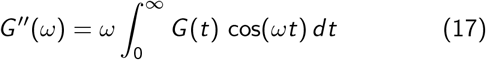

### Characterization of local geometries

In polymer physics, the correlation length of a mesh of polymers, *ξ*, can describe the entanglement spacing that arises from intermolecular interactions [83]. The general form follows:

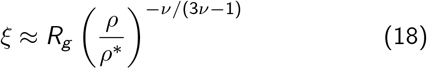

where, we define *ρ*^∗^ to represent an overlap concentration or volume occupied by a polymer chain—or in other words, the transition from the dilute to the semi-dilute regime, marking the beginning of chain interpenetration. However, condensates are not in the semi-dilute regime [84], and the *apriori* criterion of mesh isotropy prevents a characterization of heterogeneity. Therefore, we quantify local mesh environments of entangled IDPs using a radius cut-off criterion. Specifically, we look at the center of mass of individual IDP chains and collect a list of surrounding IDP chains within the cut-off radius and calculate their combined *ξ*. We justify our choice of cut-off radius by exploring a range of values: from single-molecule characterization to nearing system-wide sweeps, and selecting the radius cutoff that represents the maximum change in *ξ* (Fig. S21). Through these changes, we produce probability distributions of *ξ* which are used to describe mesh anisotropy. We also alter the definition of *ρ*^∗^ for dense phase IDP concentrations:

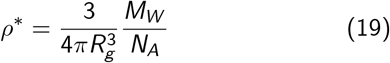

In this work, we assume that IDPs behave as ideal chains, and thus adapt a Flory scaling exponent of 1/2 [116], culminating in the form:

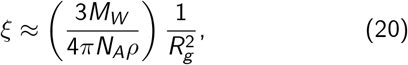

We employ the Coefficient of Variation (*CV*_*ξ*_) to characterize IDP mesh heterogeneity at different temperatures:

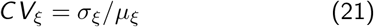

Particularly, we use *CV*_*ξ*_ to investigate how variable (*σ*_*ξ*_) the local IDP mesh geometries are relative to how their mean compaction (*µ*_*ξ*_), as this relative characterization establishes an unbiased comparison across multiple IDPs that have different chain lengths and chemistries.

### Coarse-grained simulations of dense phase systems in laminar flows

laminar shear flow simulations are set up by first equilibrating the compressed configurations of 216 proteins in the *NPT* ensemble at 0.9*T*_c_ for 50 ns, using an applied pressure of 0 atm, which allows the proteins them to achieve a density solely determined by their intermolecular interactions. Subsequently, production runs are done in the *NV T* ensemble for 100 ns. The modified Nosé–Hoover thermostat is used in these simulations via the `fix nvt/sllod` command to account for changes in simulation box size/shape and to adjusts coarse-grained bead positions. Laminar shear boundary conditions are applied via the `fix deform` command using an array of constant engineer’s strain rates. The `remap v` flag is used to adjust the velocities of coarsegrained beads once they cross periodic boundaries, consistently maintaining the generated velocity profile.

Oscillatory shear simulations are set up following the procedure of Tejedor et al. [51]. The oscillatory shear boundary conditions are applied via the fix deform command with the wiggle flag to oscillate any specified box length dimension sinusoidally with a specified amplitude and period. A total of 20 periods are simulated to obtain sufficient statistics. The period of oscillation must be at least 10 times slower than the relaxation time of the thermostat to maintain desired simulation temperatures by avoiding heat buildup from inter-IDP friction.

### Viscous dissipation and elastic character under oscillatory laminar shear flow

From oscillatory shear simulations, we obtain Lissajous curves, which are classic signatures of linear viscoelastic behavior and are directly related to hysteresis in the stress-strain rate response [7, 117]. To quantify the hysteresis of protein conformation relaxation due to shear flow, we employ Lissajous curves to qualitatively describe the degree of energy dissipation from the friction that arises as IDPs fight deformation via restoring elastic forces. The area inside the ellipse can be related to the energy dissipated per cycle. Thus, a larger area leads to greater viscous dissipation, whereas smaller areas lean towards solid-like character. The work done per cycle follows:

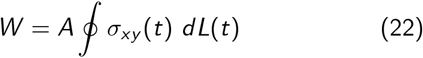

where *σ*_*xy*_ (*t*) is the ‘felt’ stress, *L*(*t*) is length of the simulation box deformed, and *A* is the constant area where the oscillatory boundary condition is enforced.

Given that the oscillatory shear simulations were conducted in the linear viscoelastic regime—deformation is directly proportional to applied strain—(Fig. S19), the shear rate and phase-shifted (*δ*) stress are obtained through:

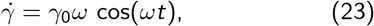

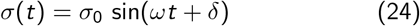

### Single chain conformational changes in laminar shear flow

We also analyze the coil-to-stretch transition of IDPs within the dense condensate environment by measuring IDP ensemble shape anisotropy, *κ*^2^, over a range of laminar shear flow rates. From the anisotropy of *R*_*g*_ tensor components from single-molecules, we can compute *κ*^2^ by first extracting the principal components, 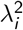, of the *Rg* tensor:

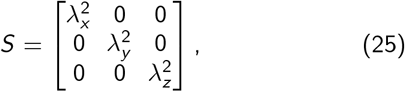

These components can then be used to calculate *κ*^2^, giving bounded values of *κ*^2^ between 0 and 1:

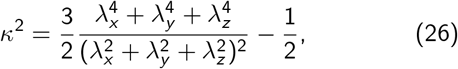

### Scalar parameters for protein sequence compositions

To attain a physical understanding of the physical characteristics of IDPs that modulate the emergent mesoscale material properties, we employ scalar parameters that describe sequence composition. First, we make use of the Sequence Hydropathy Decoration (SHD), altering the binary scale of 0 and 1 to include the Mpipi pairwise interaction strength parameter, *ε*_*ij*_ as it is a more faithful representation of amino acid residue interaction strength:

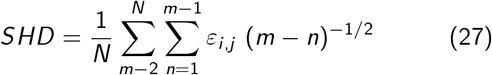

Then, to intuitively represent increasing sequence blockiness, we compute the total number of Y-S and S-Y bonds and calculate the fractional blockiness compared to sequence length, *L*, f_Blocky_ following:

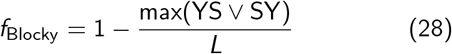

### Construction of interaction graphs

Using single static frames, we construct interaction matrices from particle position data via an energetic criterion to ensure that the interaction energy of two chains exceeds 5 *k*_B_*T* . The reason for this scalar multiplier is to select longer-lived interactions, as in Mpipi, the YY interactions are 0.42 kcal/mol which is slightly lower than *k*_B_*T* = 0.59 kcal/mol at 300 K. A 2-dimensional symmetric interaction matrix (adjacency matrix) for *N* condensed polymers, *M* = *N* × *N*, is constructed, where *M*_*ab*_ is assigned as follows:

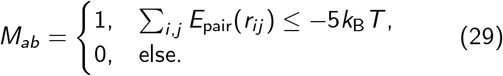

Here *i* and *j* index over each amino acid residue bead along respective protein chains *a* and *b*, and *r*_*ij*_ is the distance between monomers. Finally, graph structures are generated with the NetworkX python package [118] using binary interaction matrices as adjacency matrices *M*. Each node in a frame’s representative graph represents a single protein chain, and unweighted, undirected edges are drawn between nodes if an attractive interaction between them is observed. Associative networks are studied by finding node betweenness centrality of a node *i, C*_B_, which is found and normalized via the NetworkX `betweenness_centrality()` utility and is computed following:

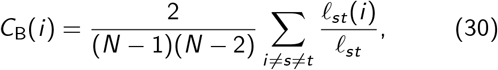

where the pair (*s, t*) enumerates over all node pairs in the graph (excluding *i* ), ℓ_*st*_ is the total number of shortest paths between *s* and *t*, and ℓ_*st*_(*i* ) is the number of shortest paths that flow through node *i* . The normalization coefficient is the inverse of the binomial coefficient for a graph with

*N* − 1 nodes, enumerating over all combinations of node pairs excluding *i* :

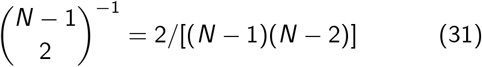

Betweenness centralities are normalized to facilitate comparison between graphs of systems of differing sizes, as the summation suggests that it is a metric that scales with the number of nodes *N*.

## ACKNOWLEDGMENTS

We thank Prof. Howard A. Stone for his invaluable insights and feedback on the manuscript. We also thank Dr. Dilimulati Aierken and Dr. Yashraj Wani for their useful comments and discussions, as well as the other members of the Joseph Group. J.A.J. also acknowledges research support from the Chan Zuckerberg Initiative DAF (an advised fund of Silicon Valley Community Foundation; grant 2023-332391) and the National Institute of General Medical Sciences of the National Institutes of Health under Award Number R35GM155259. The content is solely the responsibility of the authors and does not necessarily represent the official views of the National Institutes of Health and other sponsors. This work was also supported in part by grant NSF PHY-2309135 and the Gordon and Betty Moore Foundation Grant No. 2919.02 to the Kavli Institute for Theoretical Physics (KITP). All simulations in this work were performed using the Princeton Research Computing resources at Princeton University, which is a consortium of groups led by the Princeton Institute for Computational Science and Engineering (PICSciE) and the Office of Information Technology.

## CODE AND DATA AVAILABILITY

The data supporting the findings in this study, as well as sample simulation input and output files, are available at the Joseph Group GitHub repository: https://github.com/josephresearch/Heterogeneous_Meshwork_Analysis

