## Supplemental Information for "Biomolecular condensate viscoelasticity is dictated by the interplay between single-molecule shape memory and mesh reconfigurability"

(Dated: October 7, 2025)

#### CONTENTS

List of Figures S1

S1. Supplementary Figures S5

S2. Supplementary Methods S22
- Quantifying condensate mesh heterogeneity S22
- Quantifying condensate surface tension from slab simulations S22
- Quantifying condensate mobility through molecular diffusion S24

References S25

#### LIST OF FIGURES

S1 **Atomistic and Coarse-Grained Harmonic Bending Potentials**. These bending potentials are calculated from the probability distributions of the angles between alpha carbons of amino acids in both all-atomistic and coarse-grained resolution. . . . . S5

S2 **Correlation between radii of gyration for 17 intrinsically disordered proteins from experimental measurement and simulation for varying stiffness.** Data is shown for equilibrium bond angles of 120 **(a)**, 130, **(b)** and 140, **(c)** degrees corresponding to the harmonic angle potential used in the Hamiltonian of our coarse-grained model. The Pearson correlation coefficient, r, and deviation D are shown. . . . . S5

S3 **Wang–Frenkel potential potential curves for Y-S residue interactions**. These functions demonstrate the significant discrete differences of the cross-residue interaction strengths between tyrosine (Y) and serine (S) amino acids. . . . . S5

S4 **Phase diagrams for fully flexible and semi-flexible A1-LCD IDP condensates show agreement in predicted thermodynamic phase behavior.** . . . . . S6

S5 **Predicted phase diagrams for A1-LCD variant condensates.** Critical points are estimated from the law of coexistence densities and the law of rectilinear diameters. . . . . S6

S6 **Predicted phase diagrams for biologically relevant LCD condensates.** . . . . . S7

S7 **Predicted phase diagrams for Y-S condensates with varying hydrophobicity.** . . . . . S7

S8 **Predicted phase diagrams for Y-S condensates with varying blockiness.** . . . . . S8

S9 **Temperature-dependent shear Relaxation Moduli for A1-LCD condensates.** These $G(t)$ curves are constructed from the Green–Kubo formalism for isotropic geometry systems. Early time $G(t)$ represent relaxation mechanisms from intramolecular rearrangements. Later time $G(t)$ represent intermolecular relaxation mechanisms from friction in the confined condensate environment. These later times $G(t)$ are fitted to Maxwell modes to reduce noise at the tail-end. The increasing temperatures show relaxation changes towards more liquid like descriptions. . . . . S8

S10 **Temperature-dependent shear Relaxation Moduli for biologically relevant LCD condensates.** . . . . . S9

S11 **Temperature-dependent shear Relaxation Moduli for Y-S condensates with varying hydrophobicity.** S9

S12 **Temperature-dependent shear Relaxation Moduli for Y-S condensates with varying blockiness.** Extreme blockiness sequences like $(S_{24}Y_6)_5$ and $(S_{38}Y_{12})_3$ produce condensates that behave as glassy solids. S10

S13 **Arrhenius relationship between viscosity and temperature for experimental and simulation results of A1-LCD condensates.** The trends between A1-LCD variant condensates are shown to be relatively conserved in experimental **(a)** and simulation **(b)** results. . . . . S10

S14 **Arrhenius relationship between viscosity and normalized temperature for all studied condensates.** . . S11
S15 **Temperature-dependent loss tangents for A1-LCD variant condensates.** Different relaxation modes over frequency space are represented by the minima of $G''/G'$. Frequency 'plateaus' of $G''/G'$ are highlighted for each temperature which are used to construct the extent of elasticity analytical framework. . . . . . . . . . . . S12
S16 **Temperature-dependent loss tangents for biologically relevant LCD condensates.** . . . . . . . . . . . . . . . . S12
S17 **Temperature-dependent loss tangents for Y-S condensates with varying hydrophobicity.** . . . . . . . . . . S13
S18 **Temperature-dependent loss tangents for Y-S condensates with varying blockiness.** The shear relaxation modulus, $G(t)$, for the last variant, $(S_{38}Y_{12})_3$, has a large discontinuity, which leads to very noisy Fourier Transforms and thus, the misshaped Loss Tangent. . . . . . . . . . . . . . . . . . . . . . . . . . . . . . . . . . . . . . . . . . S13
S19 **Linear Viscoelastic Regime for A1-LCD condensates**. Oscillatory shear simulations were conducted at $0.9T_c$ using an array of amplitudes of deformation to ensure that the applied stress is proportional to the 'felt' stress. . . . . . . . . . . . . . . . . . . . . . . . . . . . . . . . . . . . . . . . . . . . . . . . . . . . . . . . . . . . . . . . . . . . . . . . . . . S14
S20 **Additional Lissajous plots for A1-LCD variants, biologically relevant LCDs and Y-S sequences with varying blockiness.** Lissajous plots for the biologically relevant LCDs **(a)**, **(b)** A1-LCD variants, and **(c)** Y-S sequences with varying blockiness are shown. The area integral quantifies dissipative work, $W$, which is shown here also. . . . . . . . . . . . . . . . . . . . . . . . . . . . . . . . . . . . . . . . . . . . . . . . . . . . . . . . . . . . . . . . . . . . . . S14
S21 **Characterization framework of entanglement spacing using various cut-off radius criteria.** **(a)** Snapshot of a condensate displaying the selected IDP in red, with surrounding mesh in blue. **(b–e)** The ensemble averaged entanglement spacing for the local meshes, $\langle\xi\rangle$, vs cut-off radius, $R_{\text{cut}}$, show that the changes in $\xi$ begin to level off at 30-40 Angstroms. **(f–i)** At very small, $R_{\text{cut}}$ the coefficient of variance of $\xi$, $CV(\xi)$ is high. This shows a dominance of finite-sample noise and intra-mesh fluctuations. On the other hand, at very large $R_{\text{cut}}$, the average is done across many meshes and the heterogeneity is lost. . . . . . . . . . S15
S22 **Probability distribution of A1-LCD variant condensates local meshes using an $R_{\text{cut}}$ of 30 Angstroms for a range of temperatures.** Two Gaussian functions are fitted to the distribution as to highlight the extended and compact meshes present in the condensate microstructure. . . . . . . . . . . . . . . . . . . . . . . . . . . . S15
S23 **Probability distribution of the biologically relevant LCDs condensates local meshes using an $R_{\text{cut}}$ of 30 Angstroms for a range of temperatures.** . . . . . . . . . . . . . . . . . . . . . . . . . . . . . . . . . . . . . . . . . . . . . . . . . S16
S24 **Probability distribution of the Y-S condensates with varying hydrophobicity local meshes using an $R_{\text{cut}}$ of 30 Angstroms for a range of temperatures.** . . . . . . . . . . . . . . . . . . . . . . . . . . . . . . . . . . . . . . . . . . . . . . S16
S25 **Probability distribution of the Y-S condensates with varying blockiness local meshes using an $R_{\text{cut}}$ of 30 Angstroms for a range of temperatures.** . . . . . . . . . . . . . . . . . . . . . . . . . . . . . . . . . . . . . . . . . . . . . . . . . S17
S26 **Temperature dependence of ensemble-averaged entanglement spacing, $\langle\xi\rangle$, for an $R_{\text{cut}}$ of 30 Angstroms for all studied condensates.** . . . . . . . . . . . . . . . . . . . . . . . . . . . . . . . . . . . . . . . . . . . . . . . S17
S27 **Correlation between $\langle\xi\rangle$ and $G''/G'$ for all studied condensates.** . . . . . . . . . . . . . . . . . . . . . . . . . . . . . . . S17
S28 **Mesh Reconfiguration Lifetimes for LCD condensates using different population thresholds** We find that results are qualitatively insensitive to modest variations of the population threshold. . . . . . . . . . . . . . . . . S18
S29 **Temperature-dependent probability distributions of betweenness centrality for A1-LCD variant condensates.** The modularity of the network topology is lost as thermal fluctuations increase in magnitude, leading to a diminishing population in 'cores'—or isolated cross-linked IDPs within the condensate microstructure. . . . . . . . . . . . . . . . . . . . . . . . . . . . . . . . . . . . . . . . . . . . . . . . . . . . . . . . . . . . . . . . . . . . . . . . . . . . S19
S30 **Temperature-dependent probability distributions of betweenness centrality for LCD condensates.** . S19
S31 **Temperature-dependent probability distributions of betweenness centrality for Y-S condensates with varying hydrophobicity.** . . . . . . . . . . . . . . . . . . . . . . . . . . . . . . . . . . . . . . . . . . . . . . . . . . . . . . . . . . . . . . . . S20
S32 **Temperature-dependent probability distributions of betweenness centrality for Y-S condensates with varying blockiness.** . . . . . . . . . . . . . . . . . . . . . . . . . . . . . . . . . . . . . . . . . . . . . . . . . . . . . . . . . . . . . . . . . . . . S20
S33 **Viscosities and Shear Stress calculated from non-equilibrium laminar shear flow simulations at different shear rates for all studied condensates.** Notably, all condensates are shear-thinning materials. . S21
S34 **Coil-to-stretch transition of IDPs under laminar shear flows collapse onto a single curve, which follows the behavior outlined by De Gennes for polymers in dilute solutions.** The Weissenberg number represents the shear rate scaled by the shape memory or relaxation times of the IDPs. Notably, $(S_{24}Y_6)_5$ and $(S_{38}Y_{12})_3$ IDPs demonstrate strain-hardening behavior. . . . . . . . . . . . . . . . . . . . . . . . . . . . . . . S21
S35 **Shannon Entropy of condensate entanglement spacing probability distributions** . . . . . . . . . . . . . . . . . . S22
S36 **Stress Profiles for slab simulations of DDX4 condensates at $0.9T_c$ demonstrating invariance to the number of bins used in the calculation of surface tension.** These stress profiles also show the density profile and surface tension value predicted by the Kirkwood–Buff formalism applied in LAMMPS. . . . . . . . . . S23
S37 **Temperature dependence of the surface tensions of all studied condensates.** Data is fitted to the analytical form prediction from the 3D Ising universality class. . . . . . . . . . . . . . . . . . . . . . . . . . . . . . . . . . . . S24
S38 **MSD-derived diffusion coefficients for the studied condensates over normalized temperature.** . . . . S24

S39 **Relationship between condensate bulk viscosity and bulk diffusion coefficients for A1-LCD variants over a range of temperatures.** The relationship follows the prediction of the Rouse Model. . . . . . . . . . . . . S25

Table S1: **Sequences of LCDs studied.**

| | |
|---|---|
| **WT A1-LCD** | GSMAS ASSSQ RGRSG SGNFG GGRGG GFGGN DNFGR GGNFS GRGGF GGSRG GGGYG GSGDG YNGFG NDGSN FGGGG SYNDF GNYNN QSSNF GPMKG GNFGG RSSGP YGGGG QYFAK PRNQG GYGGS SSSSS YGSGR RF |
| **allF** | GSMAS ASSSQ RGRSG SGNFG GGRGG GFGGN DNFGR GGNFS GRGGF GGSRG GGGFG GSGDG FNGFG NDGSN FGGGG SFNDF GNFNN QSSNF GPMKG GNFGG RSSGP FGGGG QFFAK PRNQG GFGGS SSSSS FGSGR RF |
| **allW** | GSMAS ASSSQ RGRSG SGNWG GGRGG GWGGN DNWGR GGNWS GRGGW GGSRG GGGWG GSGDG WNGWG NDGSN WGGGG SWNDW GNWNN QSSNW GPMKG GNWGG RSSGS GGGGG QWWAK PRNQG GWGGS SSSSS WGSGR RW |
| **allY** | GSMAS ASSSQ RGRSG SGNYG GGRGG GYGGN DNYGR GGNYS GRGGY GGSRG GGGYG GSGDG YNGYG NDGSN YGGGG SYNDY GNYNN QSSNY GPMKG GNYGG RSSGG SGGGG QYYAK PRNQG GYGGS SSSSS YGSGR RY |
| **YtoW** | GSMAS ASSSQ RGRSG SGNWG GGRGG GWGGN DNWGR GGNWS GRGGW GGSRG GGGWG GSGDG WNGWG NDGSN FGGGG SWNDW GNWNN QSSNF GPMKG GNFGG RSSGS GGGGG QWFAK PRNQG GWGGS SSSSS WGSGR RF |
| **FtoW** | GSMAS ASSSQ RGRSG SGNWG GGRGG GWGGN DNWGR GGNWS GRGGW GGSRG GGGYG GSGDG YNGWG NDGSN WGGGG SYNDW GNYNN QSSNW GPMKG GNWGG RSSGS GGGGG QWYAK PRNQG GYGGS SSSSS YGSGR RW |
| **DDX4** | MGDED WEAEI NPHMS SYVPI FEKDR YSGEN GDNFN RTPAS SSEMD DGPSR RDHFM KSGFA SGRNF GNRDA GECNK RDNTS TMGGF GVGKS FGNRG FSNSR FEDGD SSGFW RESSN DCEDN PTRNR GFSKR GGYRD GNNSE ASGPY RRGGR GSFRG CRGGF GLGSP NNDLD PDECM QRTGG LFGSR RPVLS GTGNG DTSQS RSGSG SERGG YKGLN EEVIT GSGKN SWKSE AEGGE S |
| **EWSR1** | MASTD YSTYS QAAAQ QGYSA YTAQP TQGYA QTTQA YGQQS YGTYG QPTDV SYTQA QTTAT YGQTA YATSY GQPPT GYTTP TAPQA YSQPV QGYGT GAYDT TTATV TTTQA SYAAQ SAYGT QPAYP AYGQQ PAATA PTRPQ DGNKP TETSQ PQSST GGYNQ PSLGY GQSNY SYPQV PGSYP MQPVT APPSY PPTSY SSTQP TSYDQ SSYSQ QNTYG QPSSY GQQSS YGQQS SYGQQ PPTSY PPQTG SYSQA PSQYS QQSSS YGQQS SFRQD HPSSM GVYGQ |
| **FUS** | MASND YTQQA TQSYG AYPTQ PGQGY SQQSS QPYGQ QSYSG YSQST DTSGY GQSSY SSYGQ SQNTG YGTQS TPQGY GSTGG YGSSQ SSQSS YGQQS SYPGY GQQPA PSSTS GSYGS SSQSS SYGQP QSGSY SQQPS YGGQQ QSYGQ QQSYN PPQGY GQQNQ YNS |
| **LAF1-RGG** | MESNQ SNNGG SGNAA LNRGG RYVPP HLRGG DGGAA AAASA GGDDR RGGAG GGGYR RGGGN SGGGG GGGYD RGYND NRDDR DNRGG SGGYG RDRNY EDRGY NGGGG GGGNR GYNNN RGGGG GGYNR QDRGD GGSSN FSRGG YNNRD EGSDN RGSGR SYNND RRDNG GDG |
| **TDP-43** | EPKHN SNRQL ERSGR FGGNP GGFGN QGGFG NSRGG GAGLG NNQGS NMGGG MNFGA FSINP AMMAA AQAAL QSSWG MMGML ASQQN QSGPS GNNQN QGNMQ REPNQ AFGSG NNSYS GSNSG AAIGW GSASN AGSGS GFNGG FGSSM DSKSS GWGM |
| **TIA1** | MINPV QQQNQ IGYPQ PYGQW GQWYG NAQQI GQYMP NGWQV PAYGM YGQAW NQQGF NQTQS SAPWM GPNYG VQPPQ GQNGS MLPNQ PSGYR VAGYE TN |

### S1. SUPPLEMENTARY FIGURES

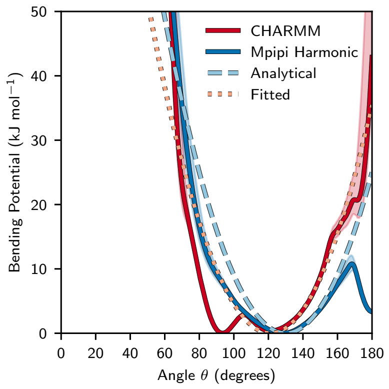


Figure S1. **Atomistic and Coarse-Grained Harmonic Bending Potentials**. These bending potentials are calculated from the probability distributions of the angles between alpha carbons of amino acids in both all-atomistic and coarse-grained resolution.

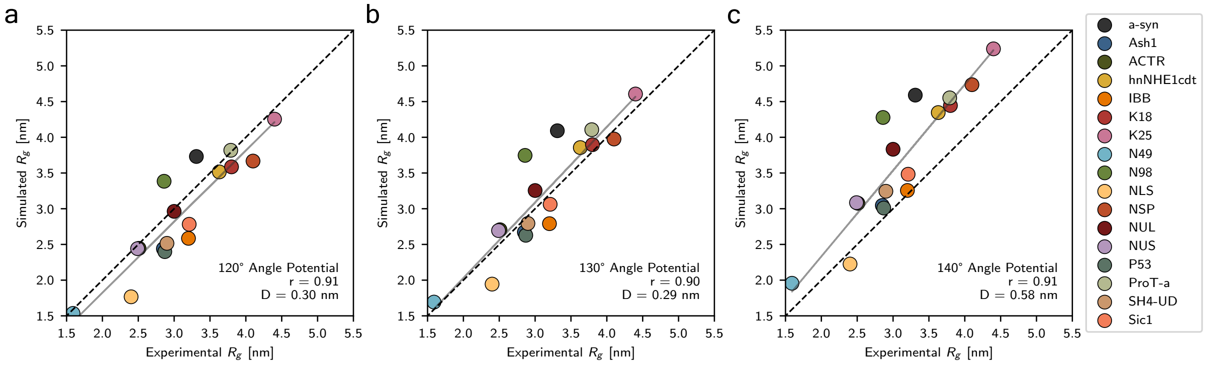


Figure S2. **Correlation between radii of gyration for 17 intrinsically disordered proteins from experimental measurement and simulation for varying stiffness.** Data is shown for equilibrium bond angles of 120 **(a)**, 130, **(b)** and 140, **(c)** degrees corresponding to the harmonic angle potential used in the Hamiltonian of our coarse-grained model. The Pearson correlation coefficient, r, and deviation D are shown.

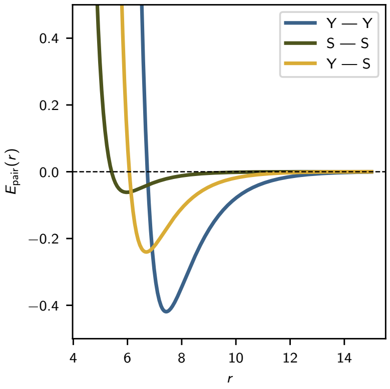


Figure S3. **Wang–Frenkel potential potential curves for Y-S residue interactions**. These functions demonstrate the significant discrete differences of the cross-residue interaction strengths between tyrosine (Y) and serine (S) amino acids.

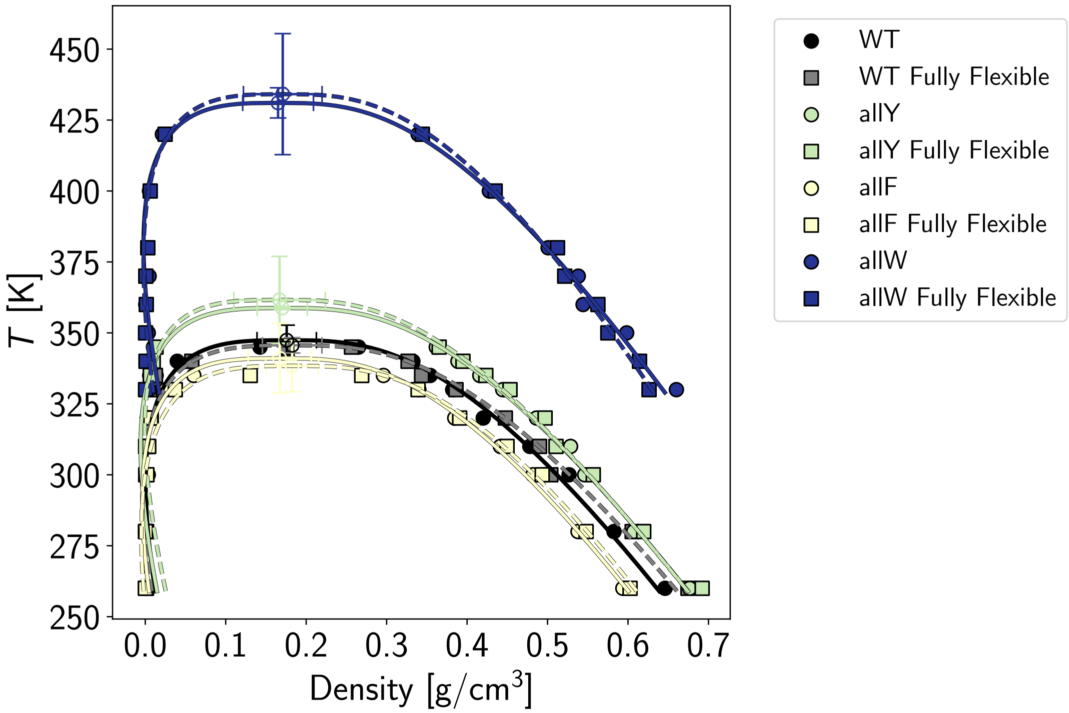


Figure S4. **Phase diagrams for fully flexible and semi-flexible A1-LCD IDP condensates show agreement in predicted thermodynamic phase behavior.**

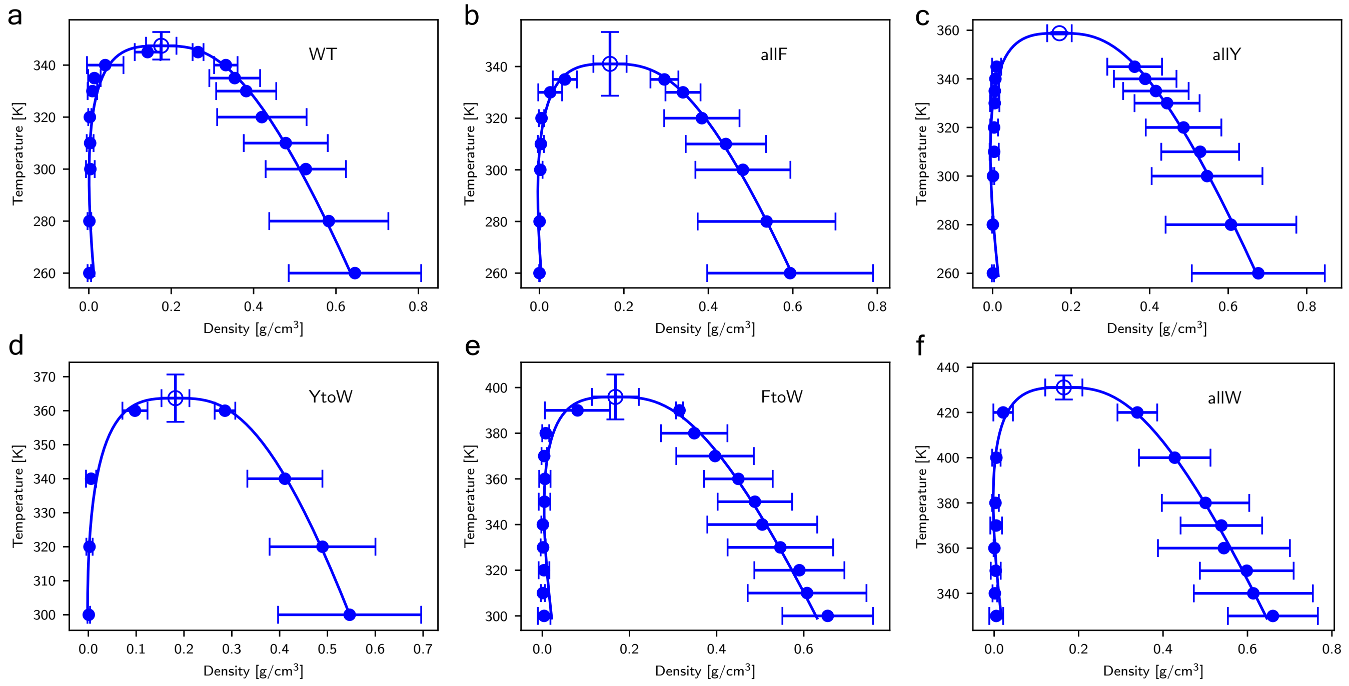


Figure S5. **Predicted phase diagrams for A1-LCD variant condensates.** Critical points are estimated from the law of coexistence densities and the law of rectilinear diameters.

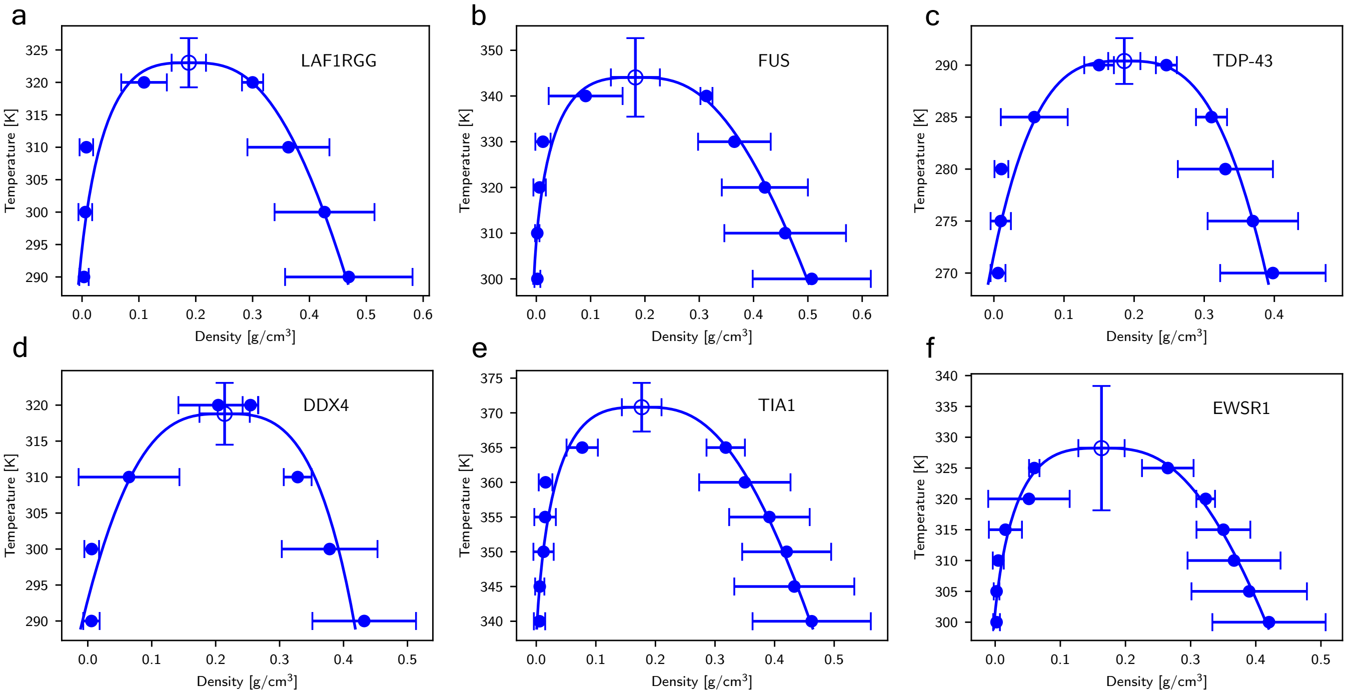


Figure S6. **Predicted phase diagrams for biologically relevant LCD condensates.**

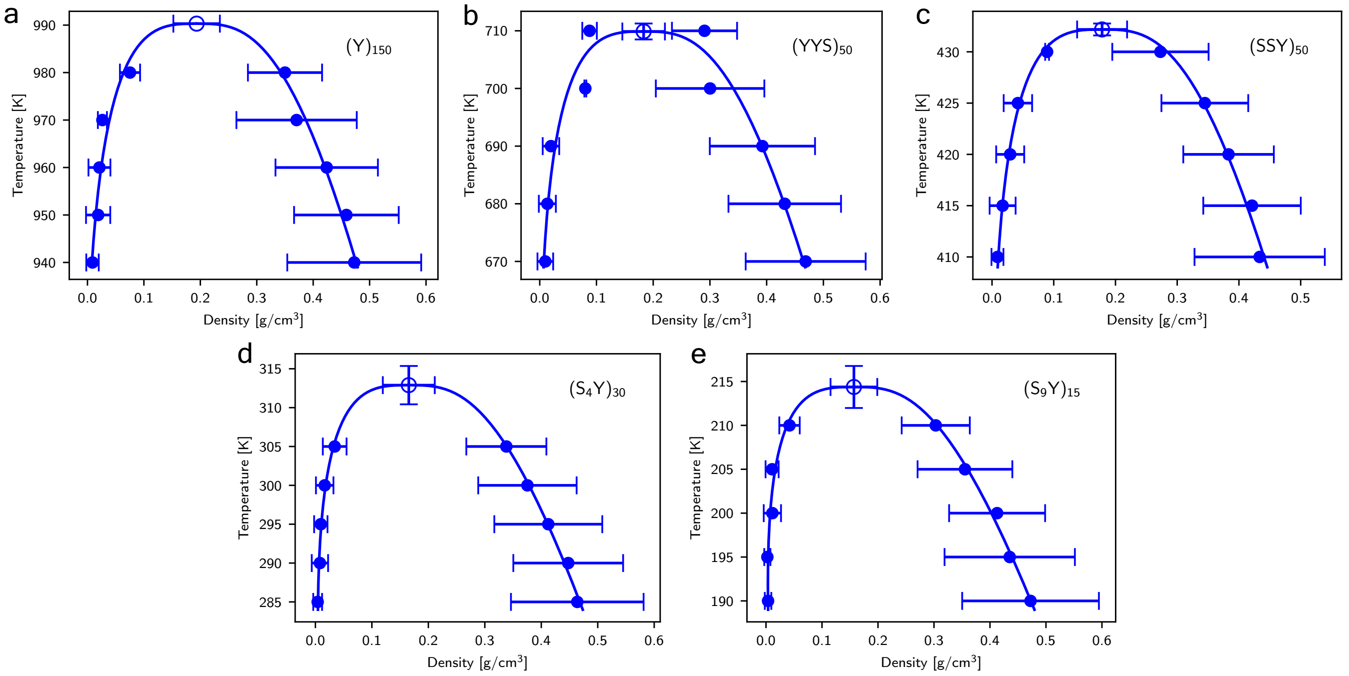


Figure S7. **Predicted phase diagrams for Y-S condensates with varying hydrophobicity.**

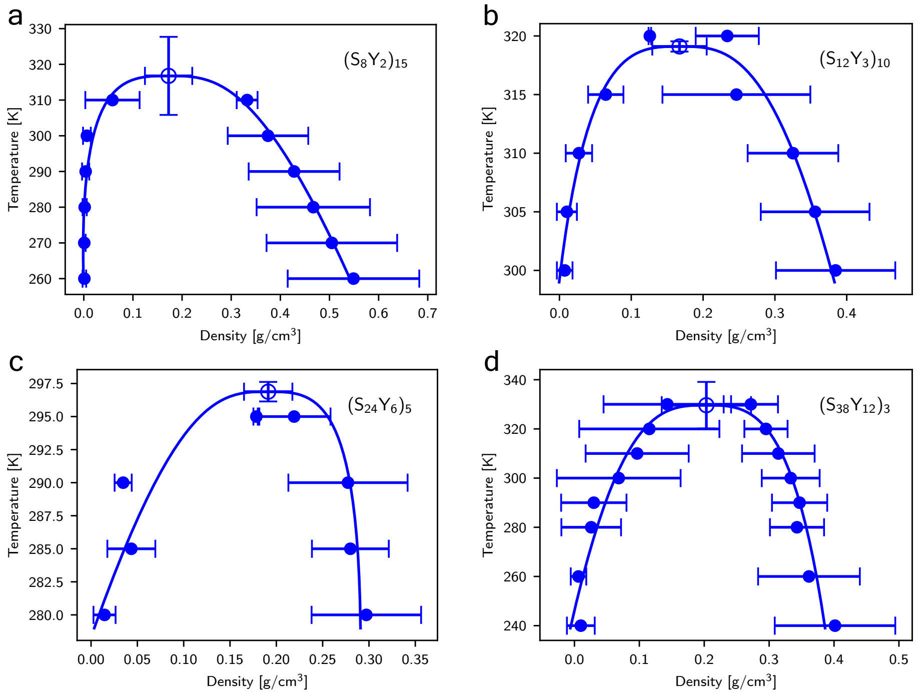


Figure S8. **Predicted phase diagrams for Y-S condensates with varying blockiness.**

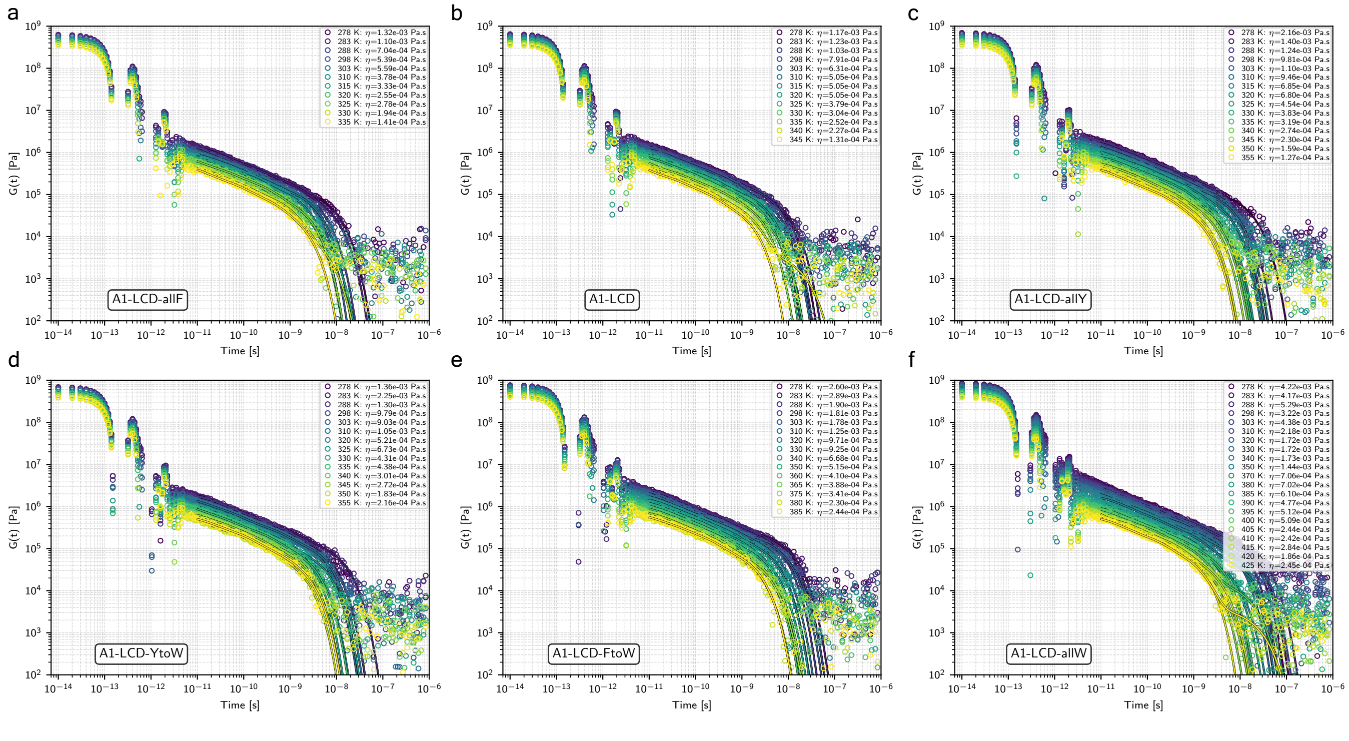


Figure S9. **Temperature-dependent shear Relaxation Moduli for A1-LCD condensates.** These $G(t)$ curves are constructed from the Green–Kubo formalism for isotropic geometry systems. Early time $G(t)$ represent relaxation mechanisms from intramolecular rearrangements. Later time $G(t)$ represent intermolecular relaxation mechanisms from friction in the confined condensate environment. These later times $G(t)$ are fitted to Maxwell modes to reduce noise at the tail-end. The increasing temperatures show relaxation changes towards more liquid like descriptions.

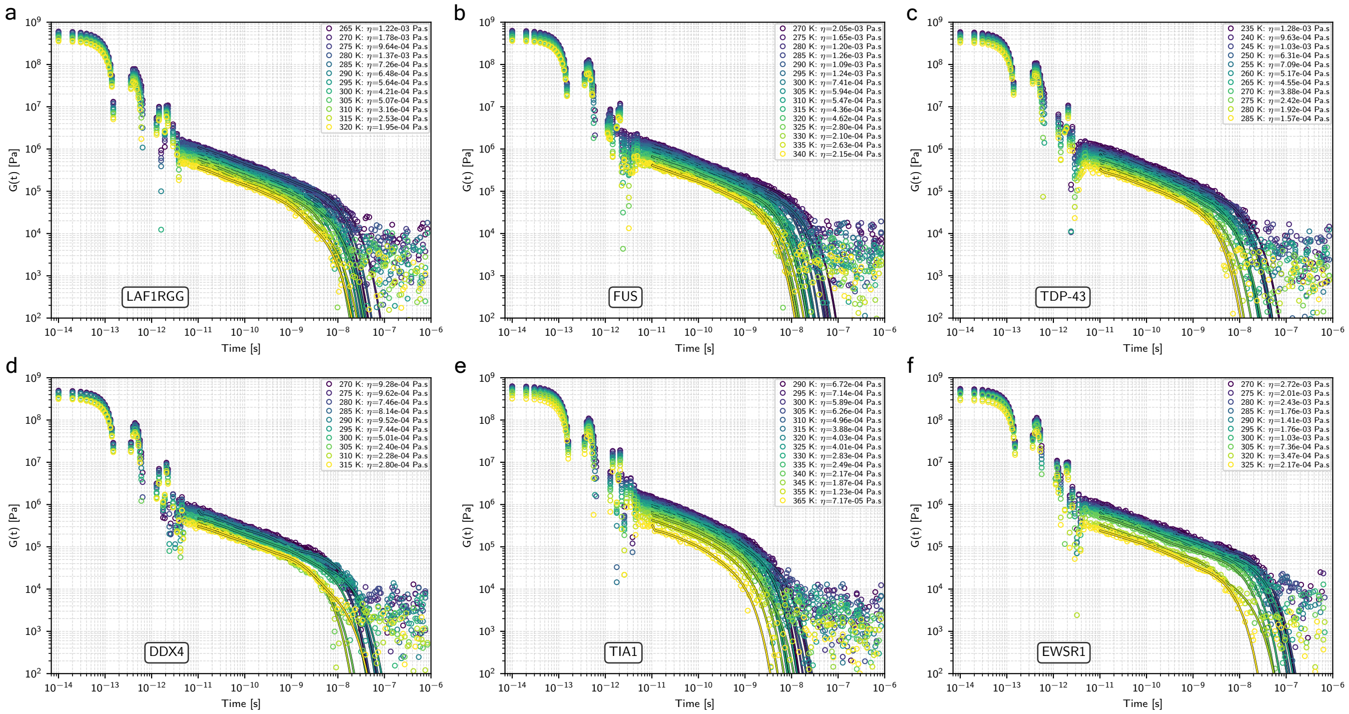


Figure S10. **Temperature-dependent shear Relaxation Moduli for biologically relevant LCD condensates.**

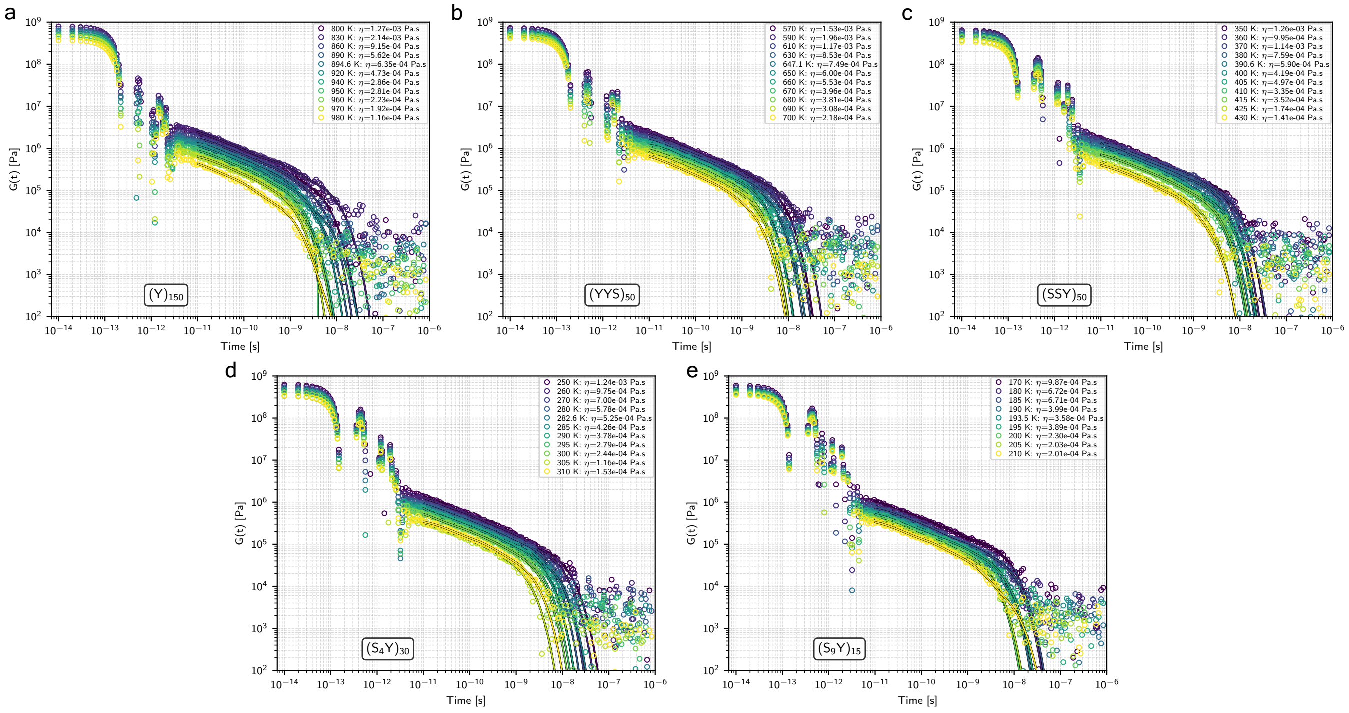


Figure S11. **Temperature-dependent shear Relaxation Moduli for Y-S condensates with varying hydrophobicity.**

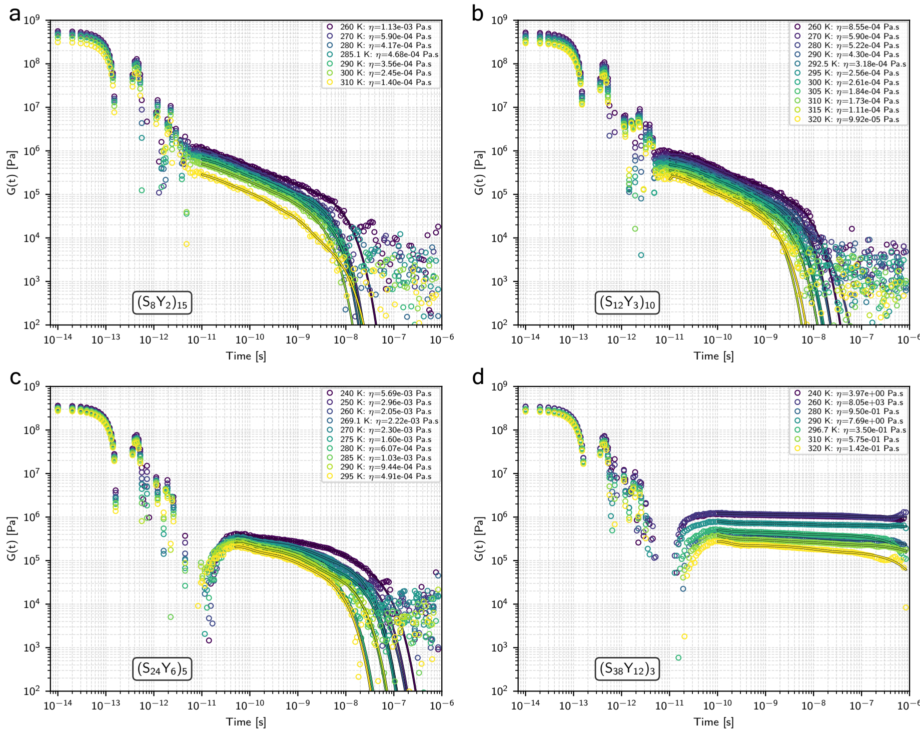


Figure S12. **Temperature-dependent shear Relaxation Moduli for Y-S condensates with varying blockiness.** Extreme blockiness sequences like $(S_{24}Y_6)_5$ and $(S_{38}Y_{12})_3$ produce condensates that behave as glassy solids.

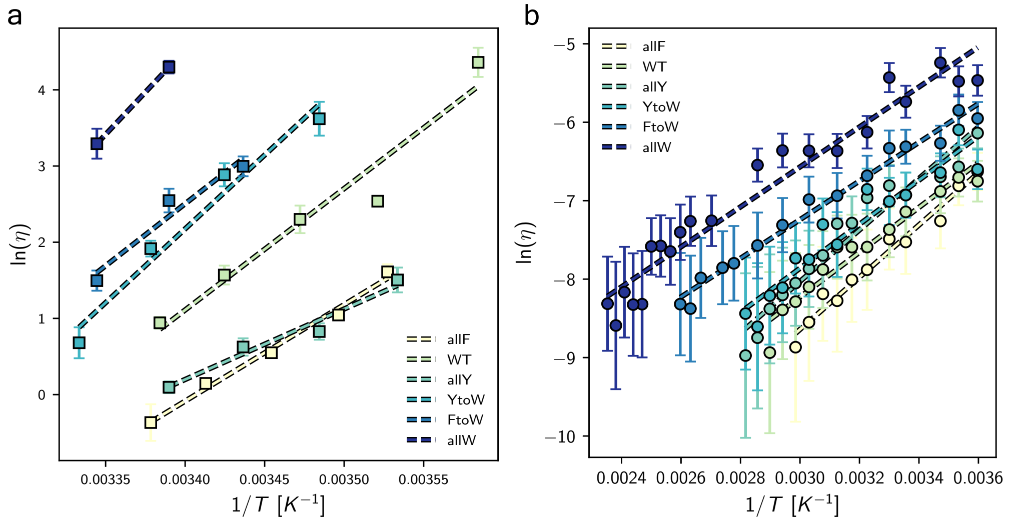


Figure S13. **Arrhenius relationship between viscosity and temperature for experimental and simulation results of A1-LCD condensates.** The trends between A1-LCD variant condensates are shown to be relatively conserved in experimental **(a)** and simulation **(b)** results.

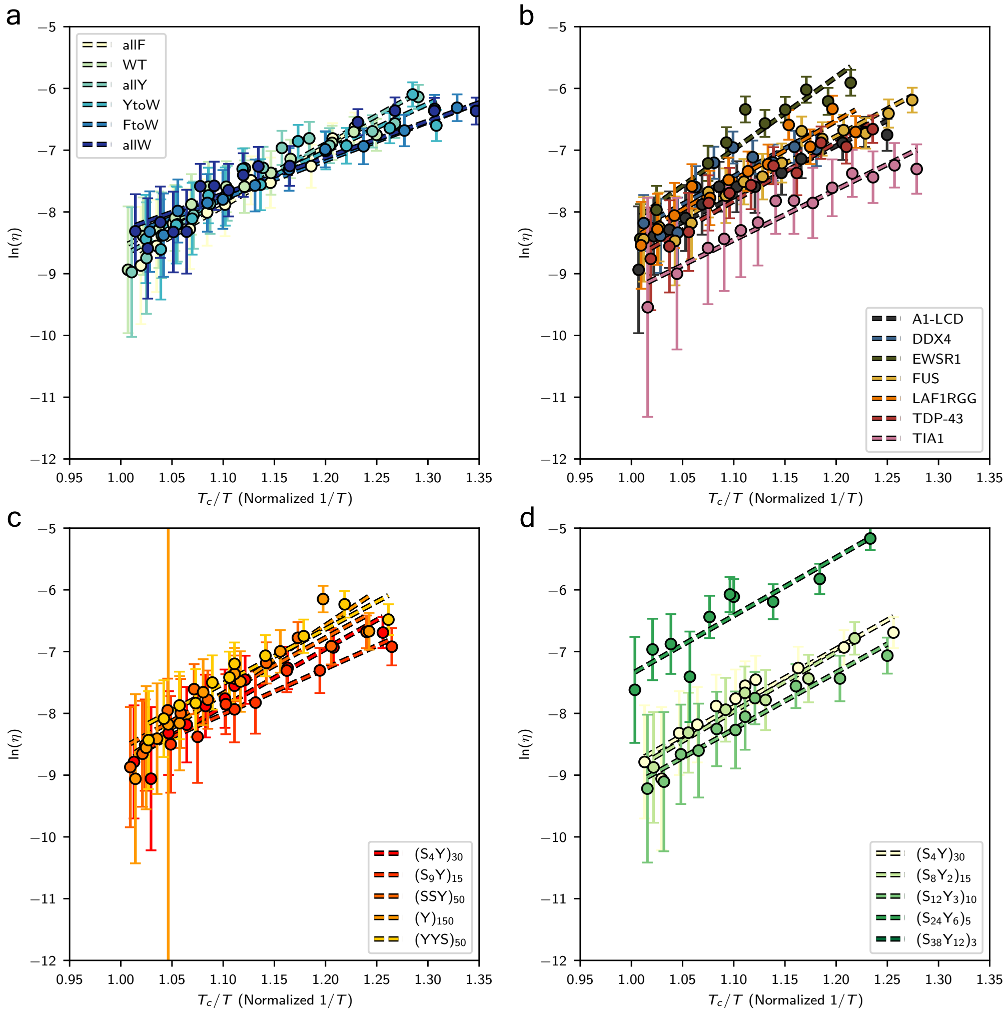


Figure S14. **Arrhenius relationship between viscosity and normalized temperature for all studied condensates.**

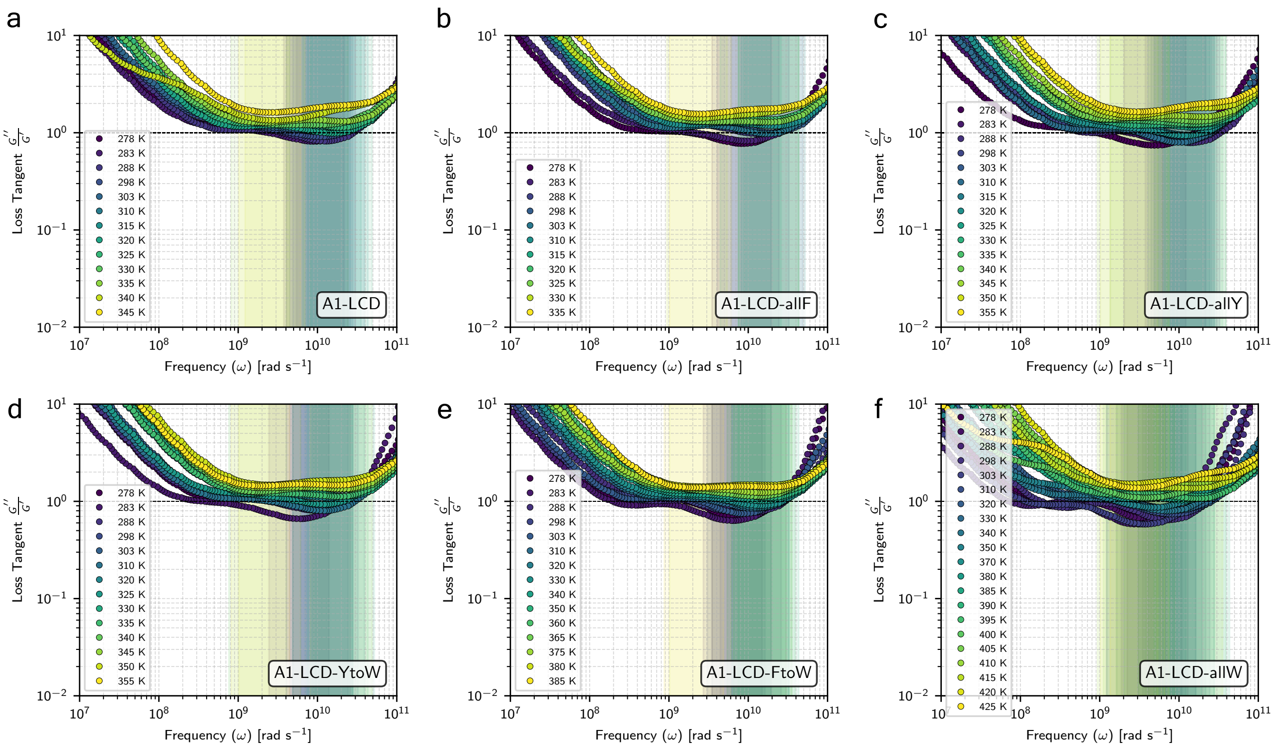


Figure S15. **Temperature-dependent loss tangents for A1-LCD variant condensates.** Different relaxation modes over frequency space are represented by the minima of $G''/G'$. Frequency 'plateaus' of $G''/G'$ are highlighted for each temperature which are used to construct the extent of elasticity analytical framework.

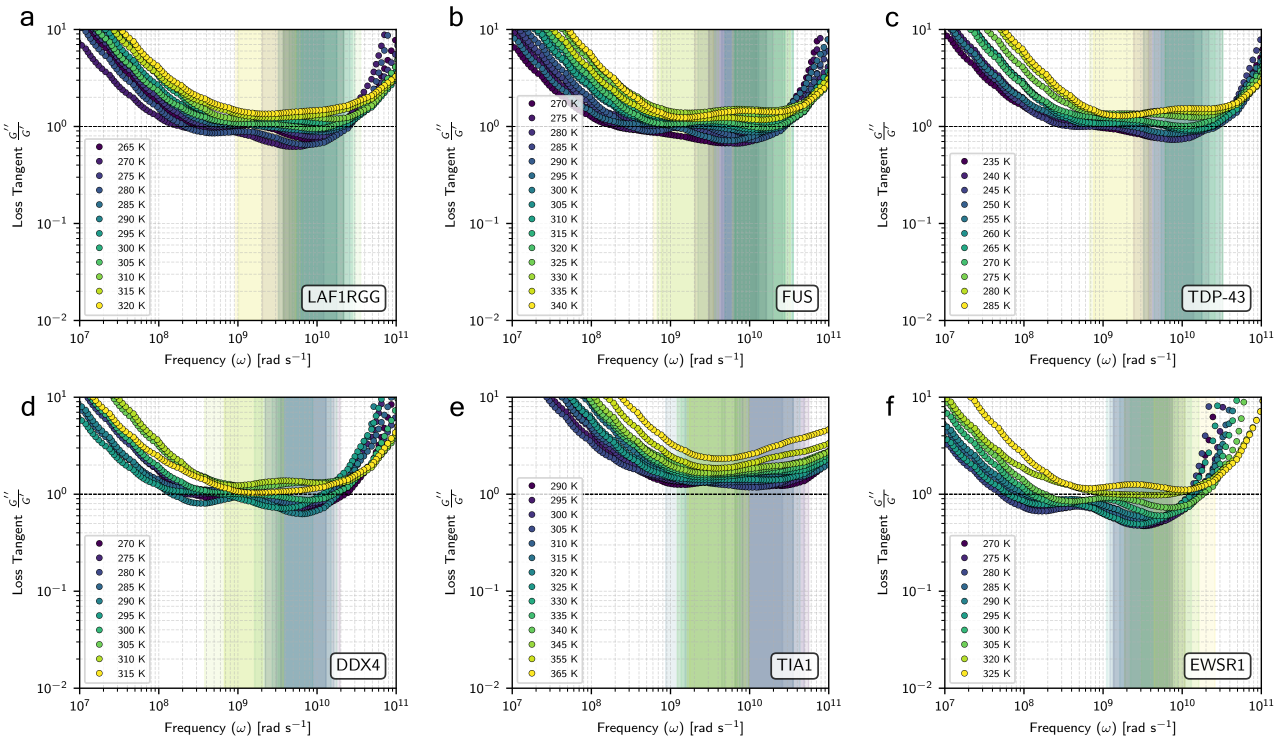


Figure S16. **Temperature-dependent loss tangents for biologically relevant LCD condensates.**

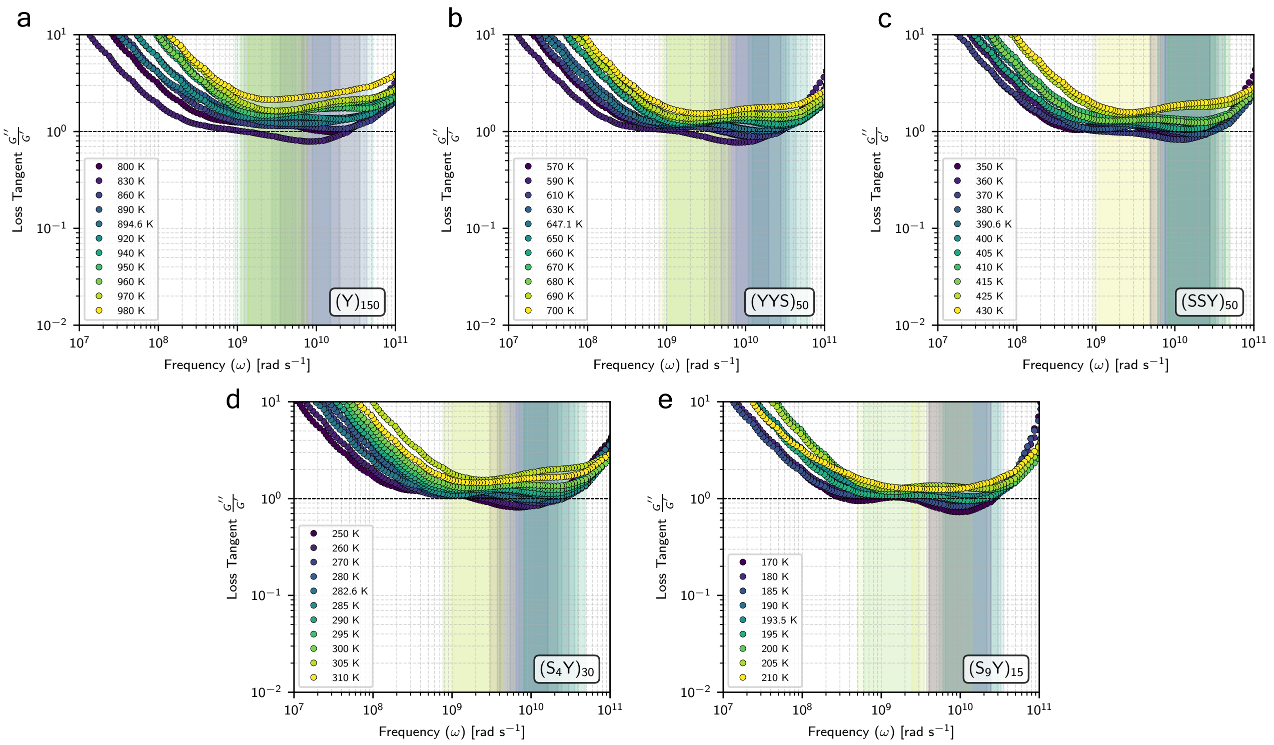


Figure S17. **Temperature-dependent loss tangents for Y-S condensates with varying hydrophobicity.**

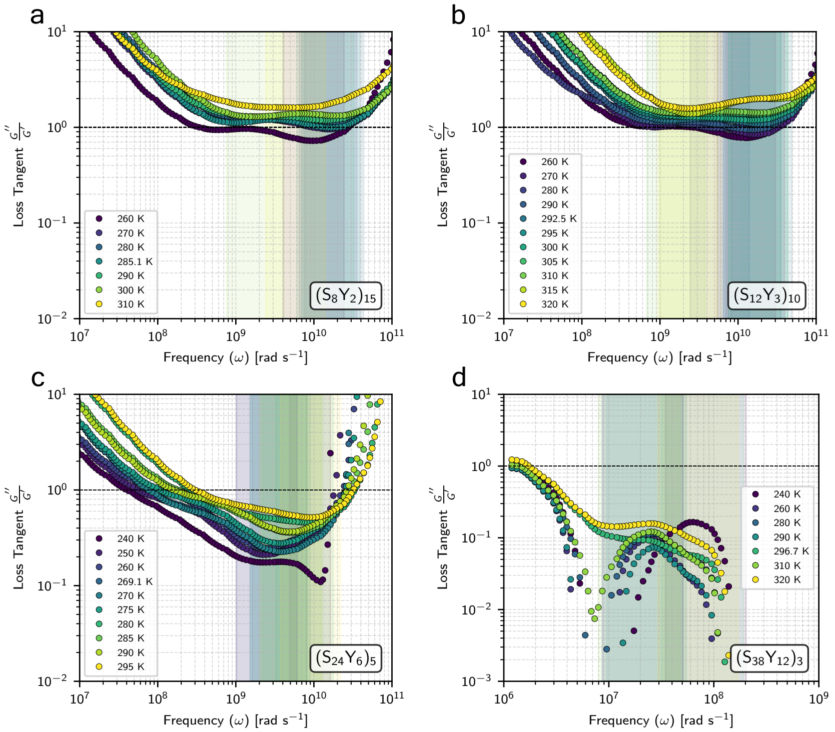


Figure S18. **Temperature-dependent loss tangents for Y-S condensates with varying blockiness.** The shear relaxation modulus, $G(t)$, for the last variant, $(S_{38}Y_{12})_3$, has a large discontinuity, which leads to very noisy Fourier Transforms and thus, the misshaped Loss Tangent.

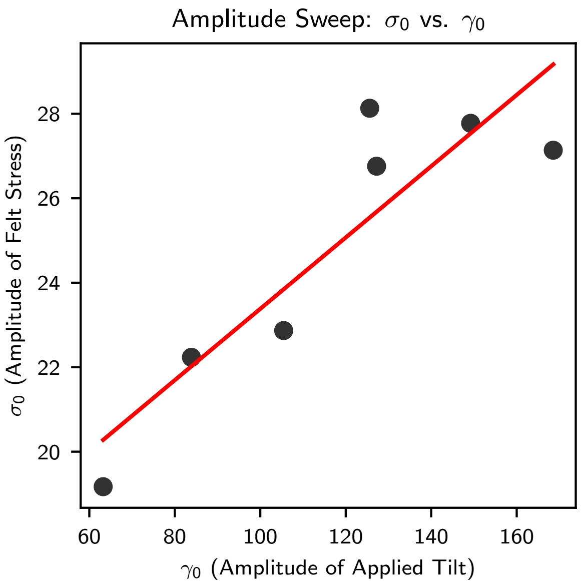


Figure S19. **Linear Viscoelastic Regime for A1-LCD condensates**. Oscillatory shear simulations were conducted at $0.9T_c$ using an array of amplitudes of deformation to ensure that the applied stress is proportional to the 'felt' stress.

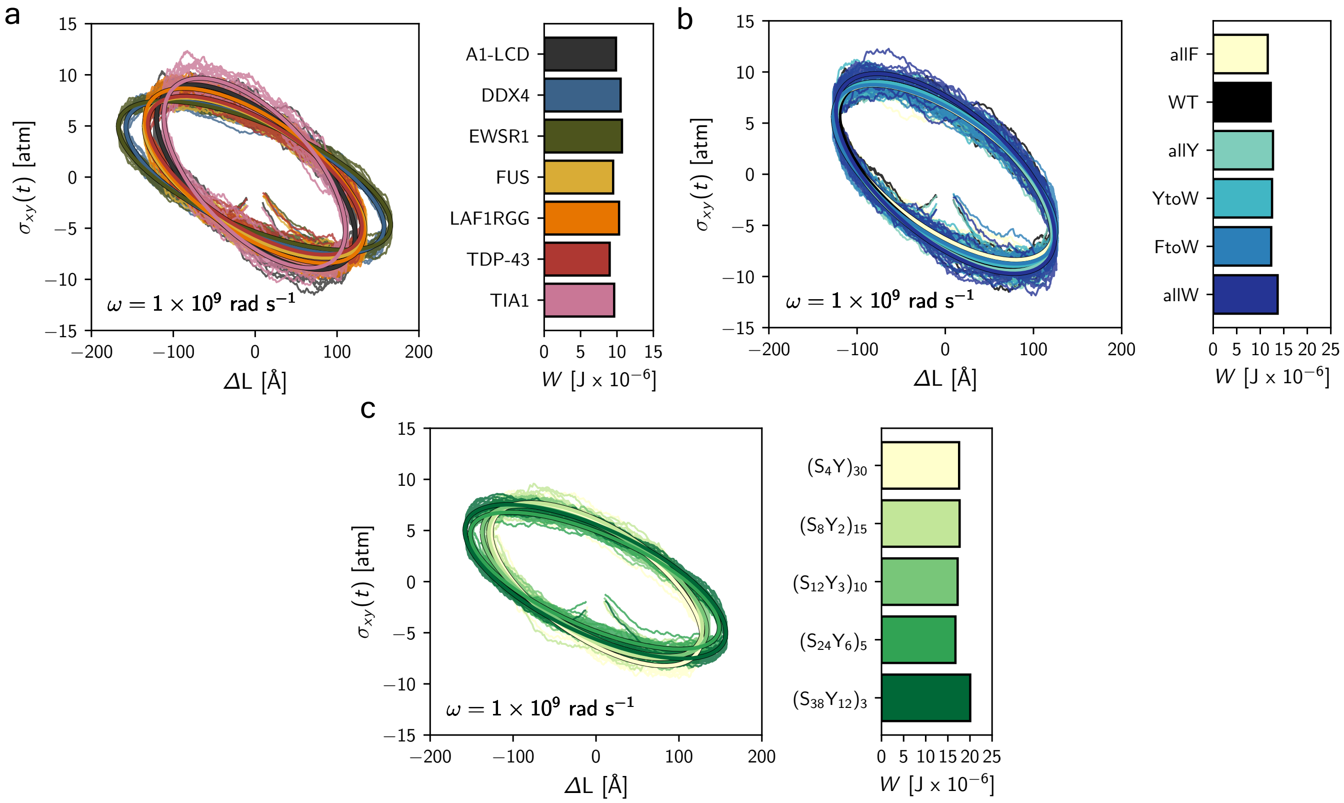


Figure S20. **Additional Lissajous plots for A1-LCD variants, biologically relevant LCDs and Y-S sequences with varying blockiness.** Lissajous plots for the biologically relevant LCDs **(a)**, **(b)** A1-LCD variants, and **(c)** Y-S sequences with varying blockiness are shown. The area integral quantifies dissipative work, $W$, which is shown here also.

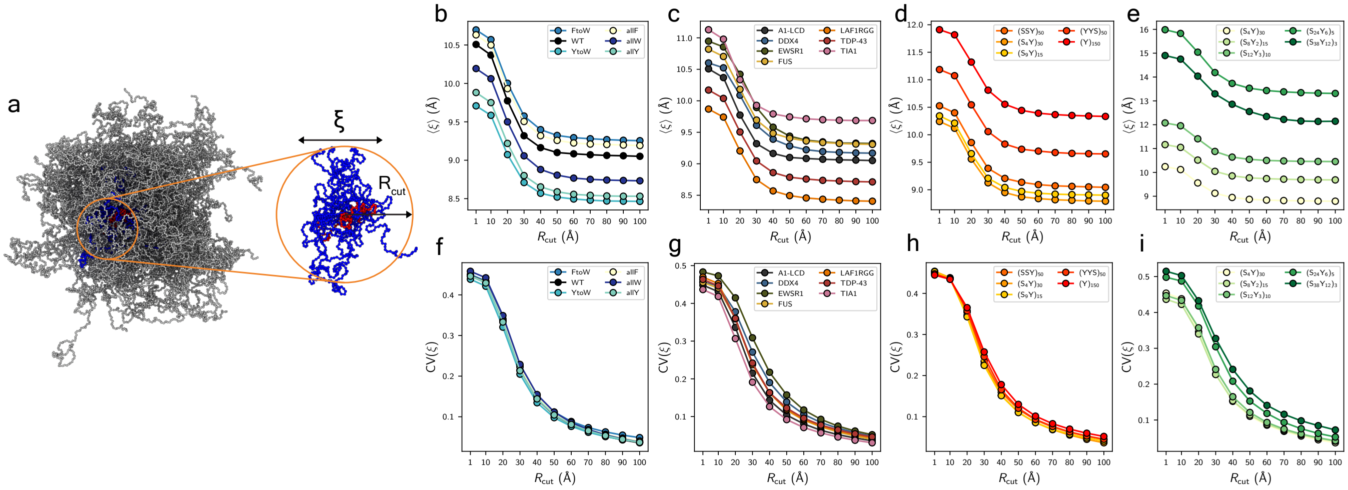


Figure S21. **Characterization framework of entanglement spacing using various cut-off radius criteria.** **(a)** Snapshot of a condensate displaying the selected IDP in red, with surrounding mesh in blue. **(b–e)** The ensemble averaged entanglement spacing for the local meshes, $\langle\xi\rangle$, vs cut-off radius, $R_{\mathrm{cut}}$, show that the changes in $\xi$ begin to level off at 30-40 Angstroms. **(f–i)** At very small, $R_{\mathrm{cut}}$ the coefficient of variance of $\xi$, $CV(\xi)$ is high. This shows a dominance of finite-sample noise and intra-mesh fluctuations. On the other hand, at very large $R_{\mathrm{cut}}$, the average is done across many meshes and the heterogeneity is lost.

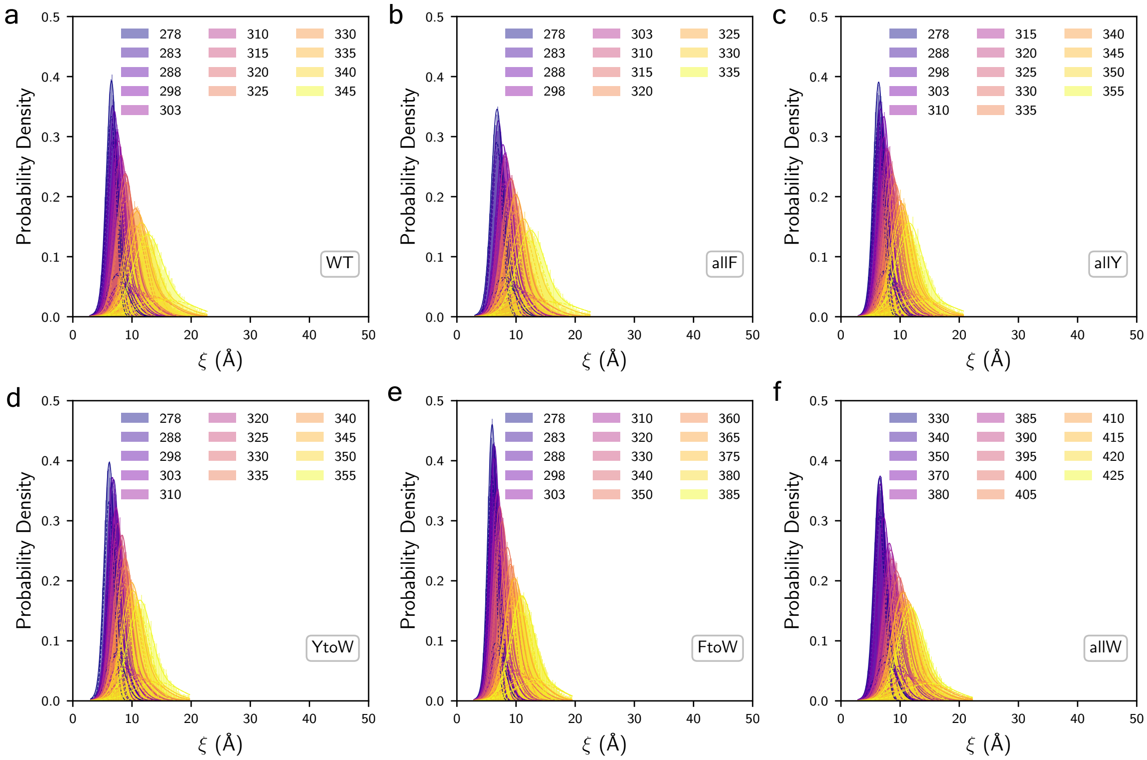


Figure S22. **Probability distribution of A1-LCD variant condensates local meshes using an $R_{\mathrm{cut}}$ of 30 Angstroms for a range of temperatures.** Two Gaussian functions are fitted to the distribution as to highlight the extended and compact meshes present in the condensate microstructure.

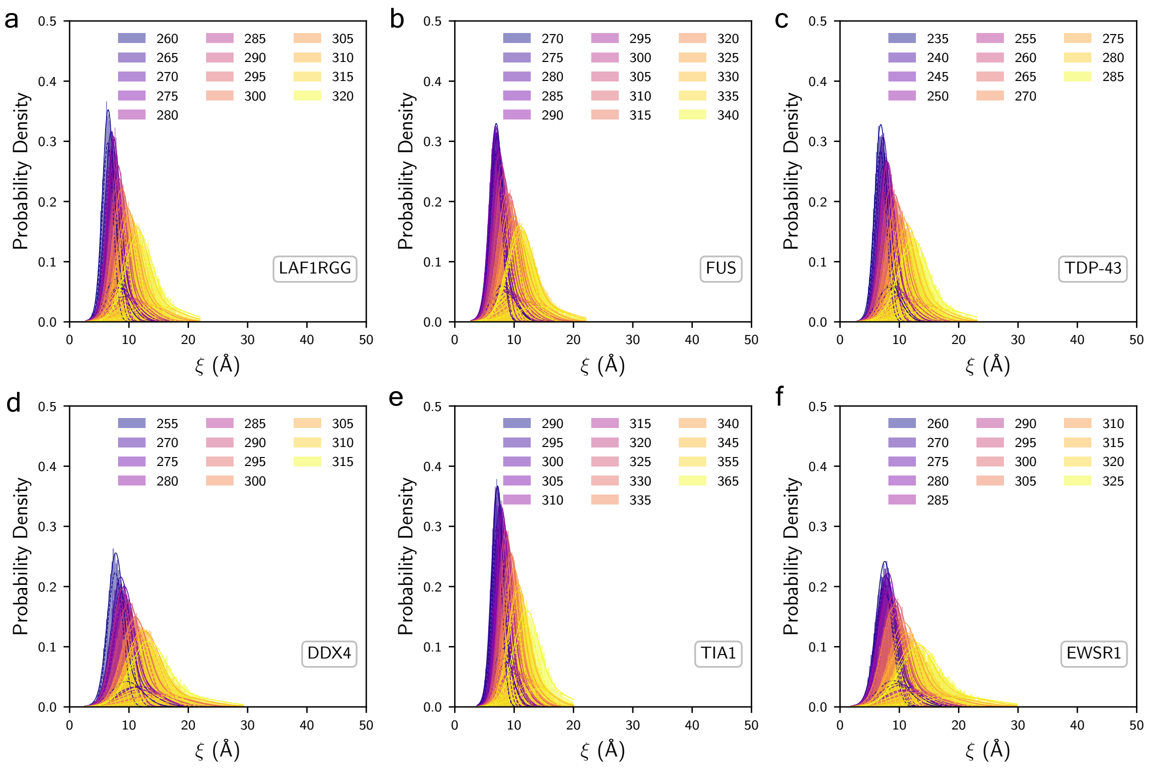


Figure S23. **Probability distribution of the biologically relevant LCDs condensates local meshes using an $R_{\mathrm{cut}}$ of 30 Angstroms for a range of temperatures.**

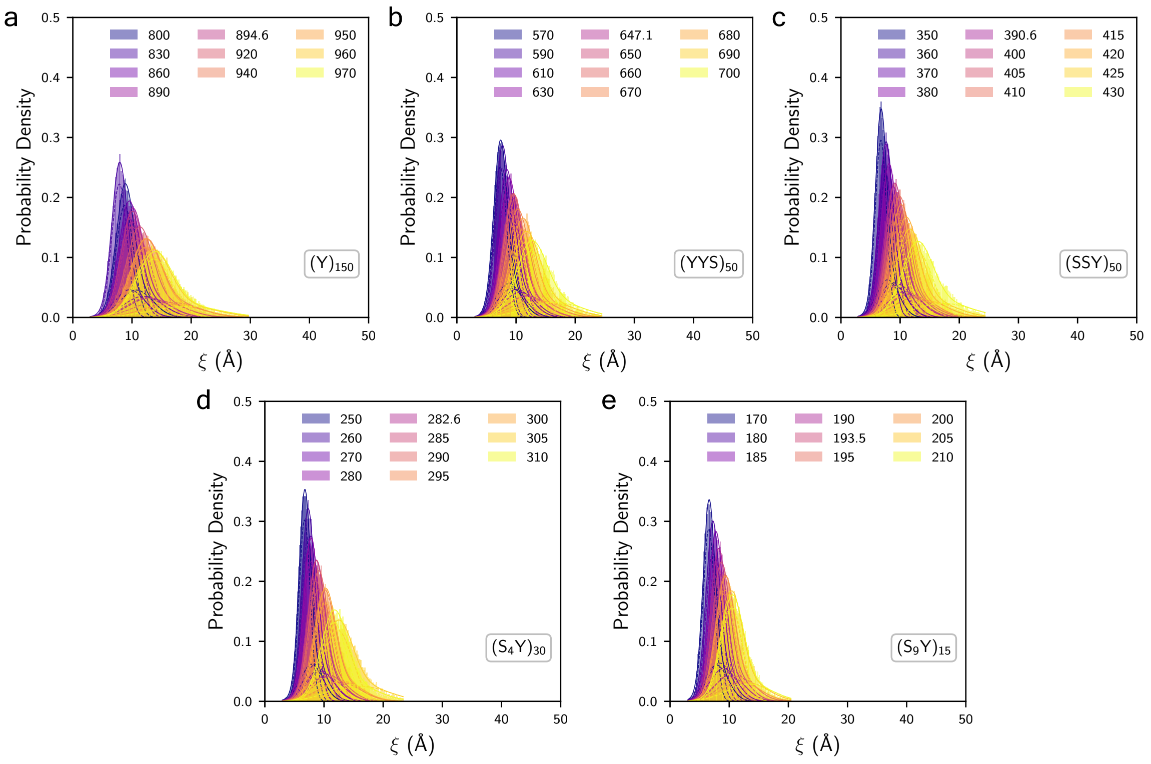


Figure S24. **Probability distribution of the Y-S condensates with varying hydrophobicity local meshes using an $R_{\mathrm{cut}}$ of 30 Angstroms for a range of temperatures.**

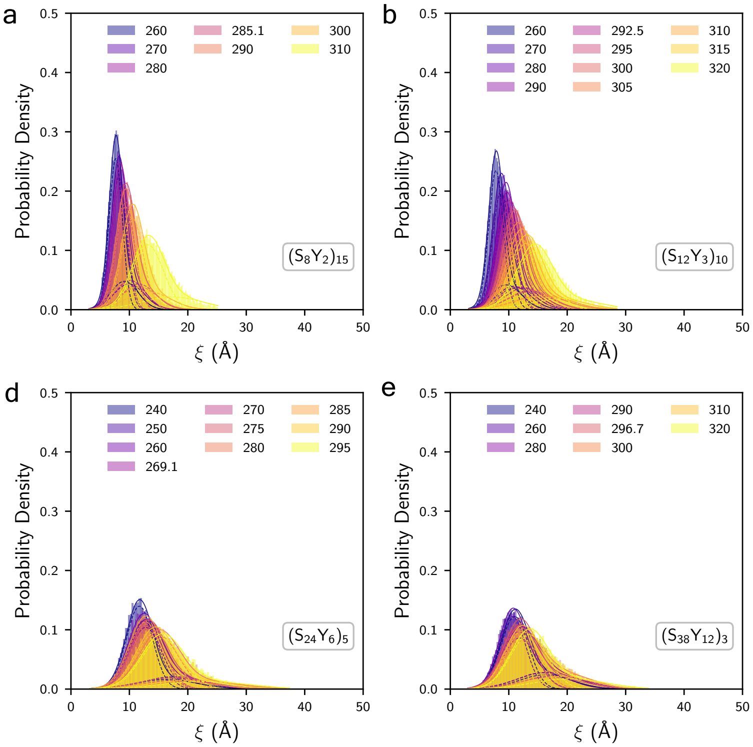


Figure S25. **Probability distribution of the Y-S condensates with varying blockiness local meshes using an $R_{\mathrm{cut}}$ of 30 Angstroms for a range of temperatures.**

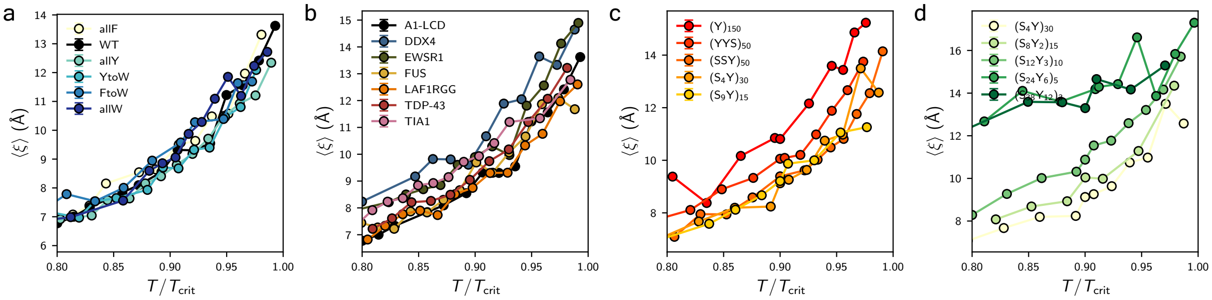


Figure S26. **Temperature dependence of ensemble-averaged entanglement spacing, $\langle \xi \rangle$, for an $R_{\mathrm{cut}}$ of 30 Angstroms for all studied condensates.**

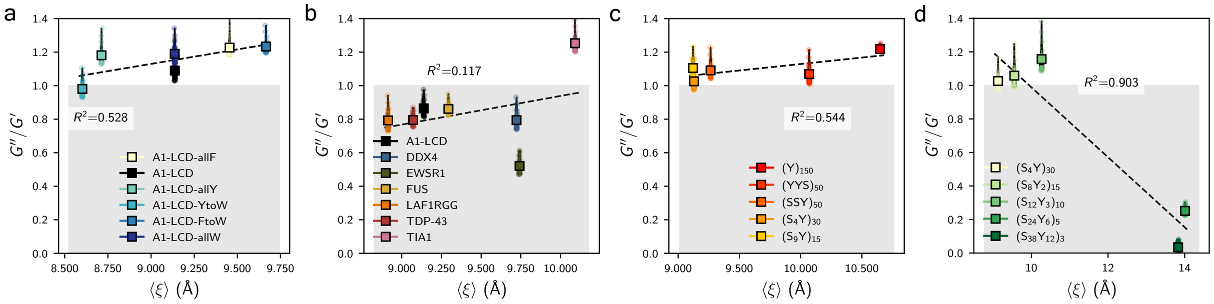


Figure S27. **Correlation between $\langle \xi \rangle$ and $G''/G'$ for all studied condensates.**

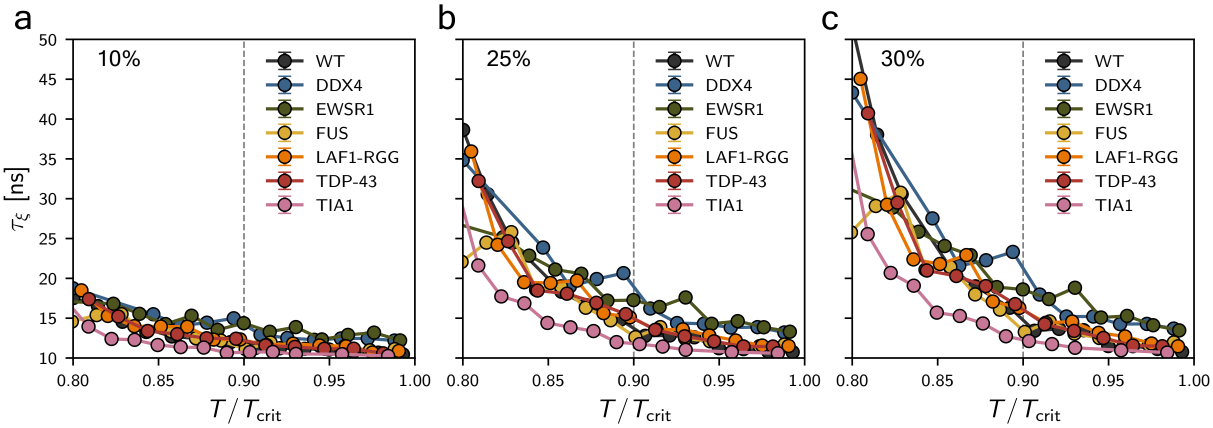


Figure S28. **Mesh Reconfiguration Lifetimes for LCD condensates using different population thresholds** We find that results are qualitatively insensitive to modest variations of the population threshold.

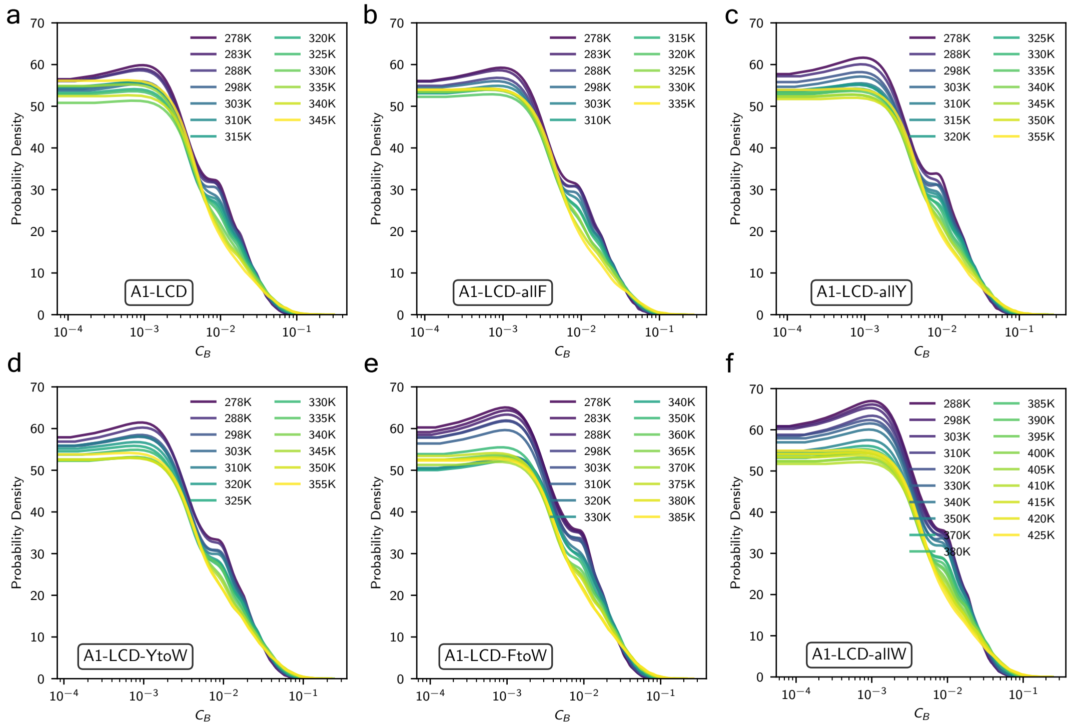


Figure S29. **Temperature-dependent probability distributions of betweenness centrality for A1-LCD variant condensates.** The modularity of the network topology is lost as thermal fluctuations increase in magnitude, leading to a diminishing population in 'cores'—or isolated cross-linked IDPs within the condensate microstructure.

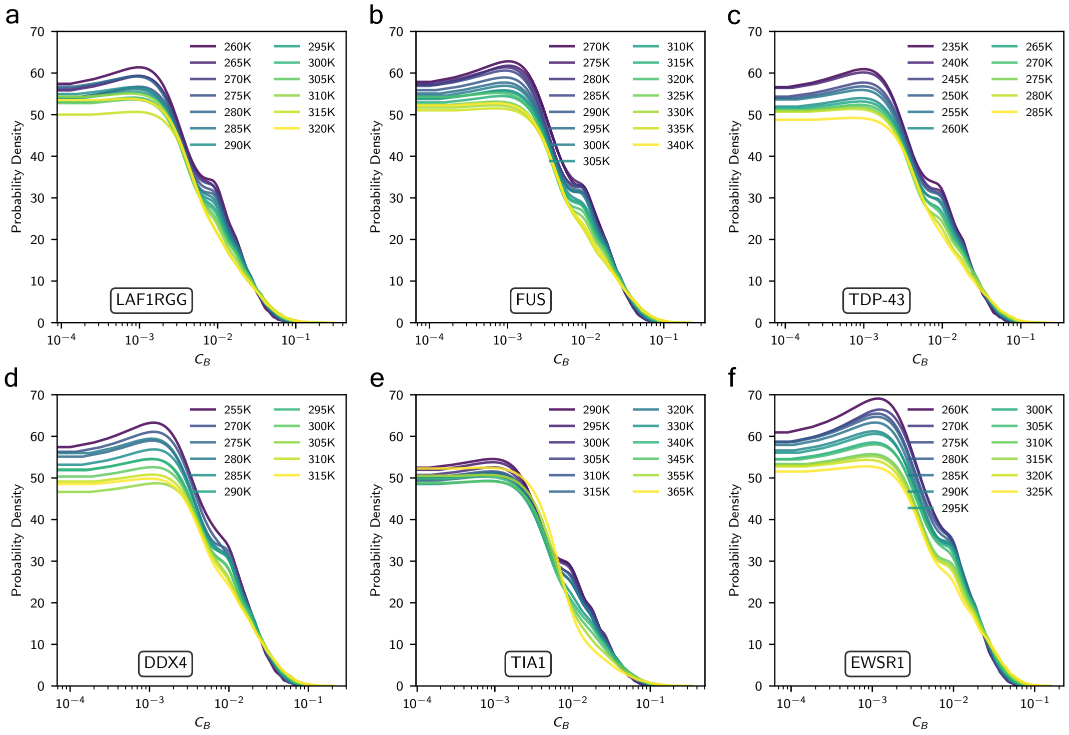


Figure S30. **Temperature-dependent probability distributions of betweenness centrality for LCD condensates.**

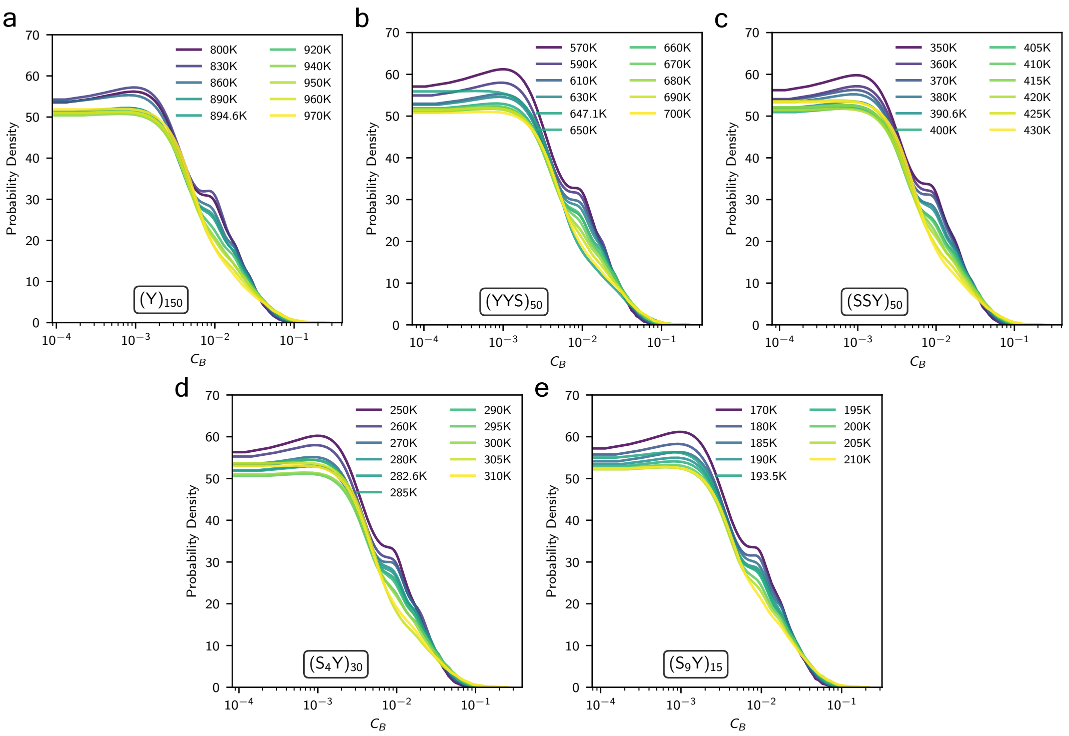


Figure S31. **Temperature-dependent probability distributions of betweenness centrality for Y-S condensates with varying hydrophobicity.**

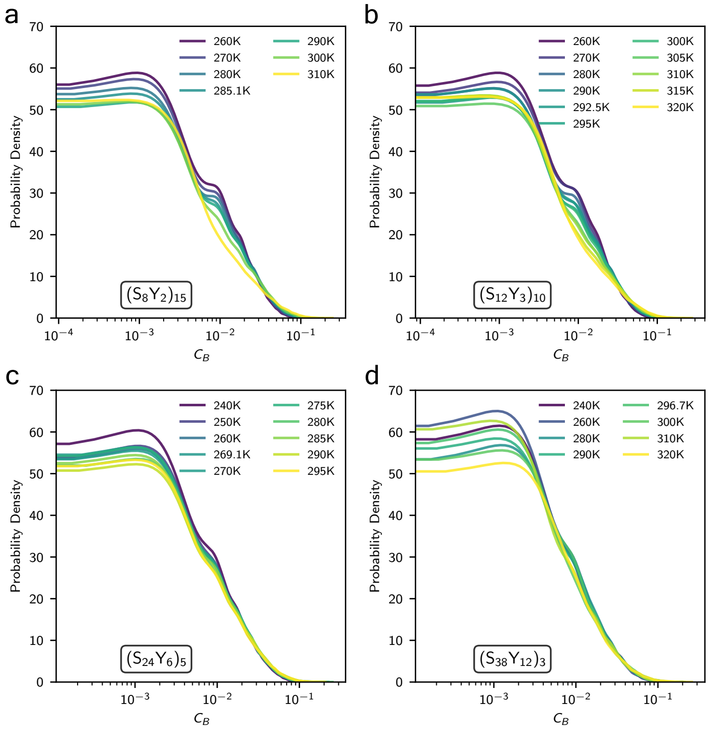


Figure S32. **Temperature-dependent probability distributions of betweenness centrality for Y-S condensates with varying blockiness.**

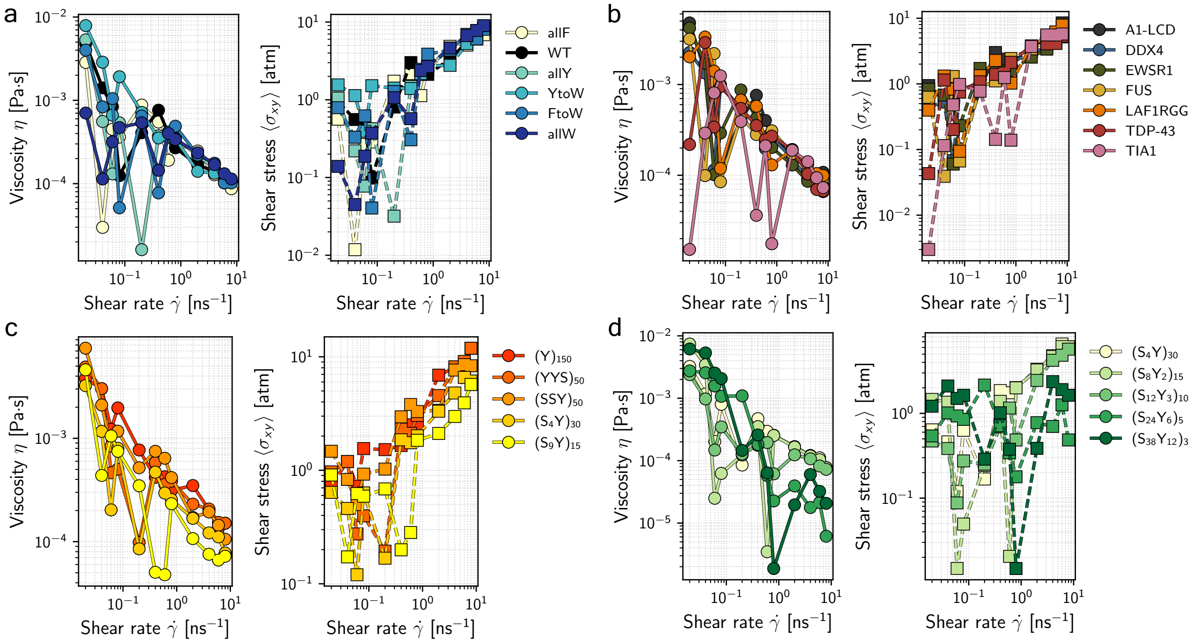


Figure S33. **Viscosities and Shear Stress calculated from non-equilibrium laminar shear flow simulations at different shear rates for all studied condensates.** Notably, all condensates are shear-thinning materials.

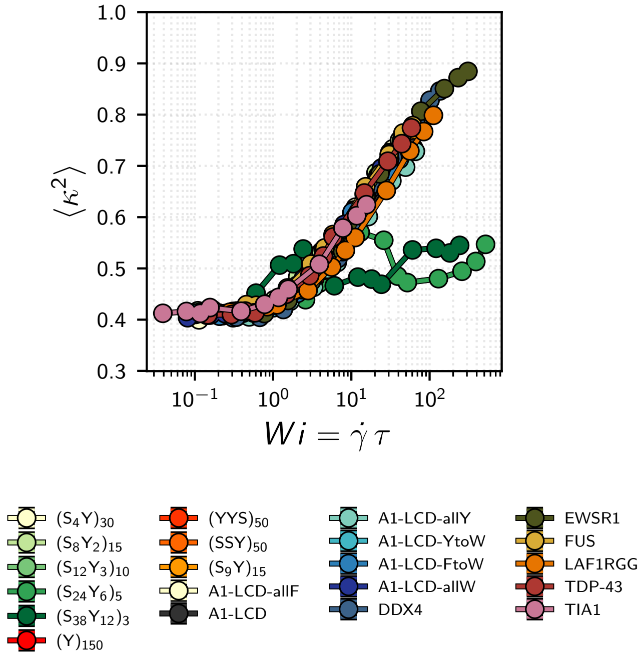


Figure S34. **Coil-to-stretch transition of IDPs under laminar shear flows collapse onto a single curve, which follows the behavior outlined by De Gennes for polymers in dilute solutions.** The Weissenberg number represents the shear rate scaled by the shape memory or relaxation times of the IDPs. Notably, $(S_{24}Y_6)_5$ and $(S_{38}Y_{12})_3$ IDPs demonstrate strain-hardening behavior.

#### S2. SUPPLEMENTARY METHODS

*Quantifying condensate mesh heterogeneity*

We also consider the **Shannon Entropy** as a descriptor of the spread of $\mathcal{P}(\xi)$. By presenting our results for this metric, we wish to reinforce the CV($\xi$) results and isolate just the shape or inequality of the distribution, where high entropy means a spread out distribution, and a more heterogeneous mesh:

$$H_{\mathrm{norm}}(X) = -\frac{1}{\log_2(k)} \sum \mathcal{P}(X) \log \mathcal{P}(X) \tag{S1}$$

here, k is the number of non-zero bins from the calculated histograms.

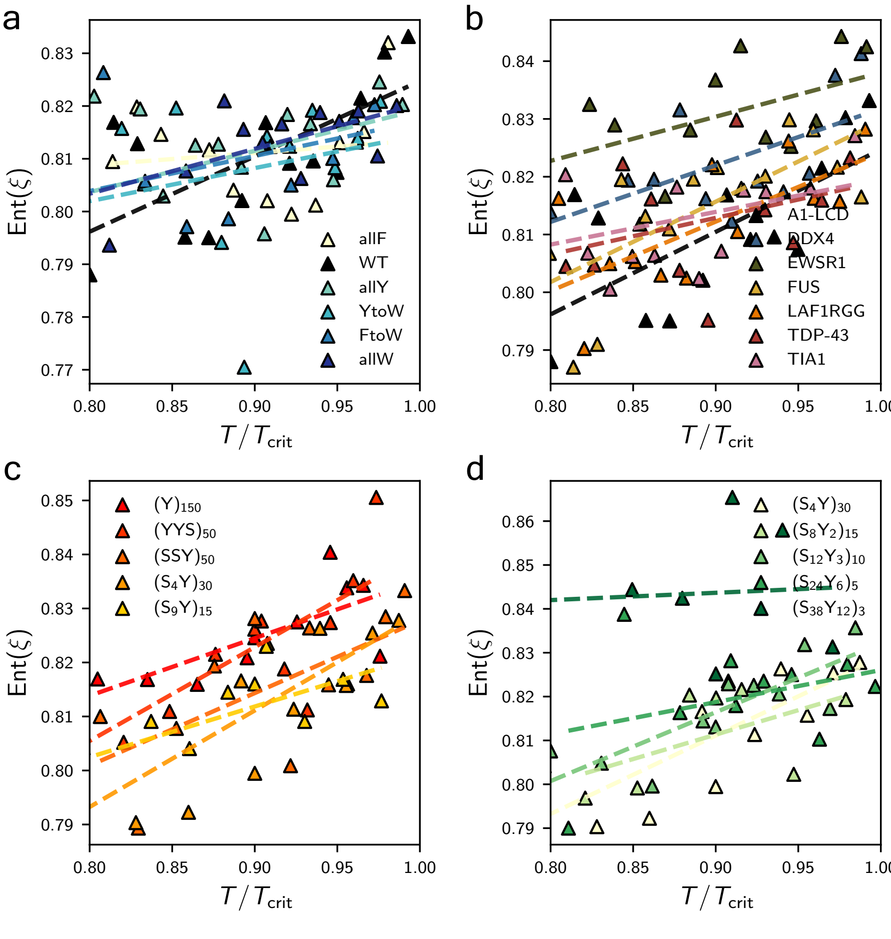


Figure S35. **Shannon Entropy of condensate entanglement spacing probability distributions**

*Quantifying condensate surface tension from slab simulations*

To assess whether our new coarse-grained model Hamiltonian altered other thermodynamic properties of condensates predicted by the original *Mpipi* model, we calculate the **surface tensions** of condensates from slab simulations. The surface tension between two phases of different densities is usually described by the Kirkwood–Buff formalism [1, 2], considering that the presence of an interface provides anisotropy to the overall pressure tensor. Thus, following the definition of mechanical equilibrium, the surface tension is given by:

$$\gamma = \frac{L_z}{2}\left[\langle\sigma_{zz}\rangle - \frac{1}{2}(\langle\sigma_{xx}\rangle + \langle\sigma_{yy}\rangle)\right] \quad \text{(S2)}$$

where, $L_z$ is the length of the slab, and $\langle\sigma_{ii}\rangle$ are the ensemble averages of the diagonal components of the pressure tensor.

However, the formalism is built upon the assumption of a sharp interface, which breaks down as a condensate's critical temperature is approached. At that limit, the interface become more diffuse and harder to define, yielding incorret surface tension estimations through the Kirkwood-Buff formalism. To overcome this difficulty, we employ a stress-profile method calculated from per-atom stresses using the virial force contribution equation [3]. This derivation is possible because anisotropy of the diagonal components of the pressure tensors are found at the dense-dilute interface:

$$\gamma = \frac{L_z}{2}\sum_{z\in\{z_L,z_R\}}\int_{z-w/2}^{z+w/2}\langle\sigma_{zz}\rangle - \tfrac{1}{2}(\langle\sigma_{xx}\rangle + \langle\sigma_{yy}\rangle)\; dz \quad \text{(S3)}$$

here, $z_L$, $z_R$ correspond to the interfaces of the dense phase in the slab geometry, with an equal thickness $w$.

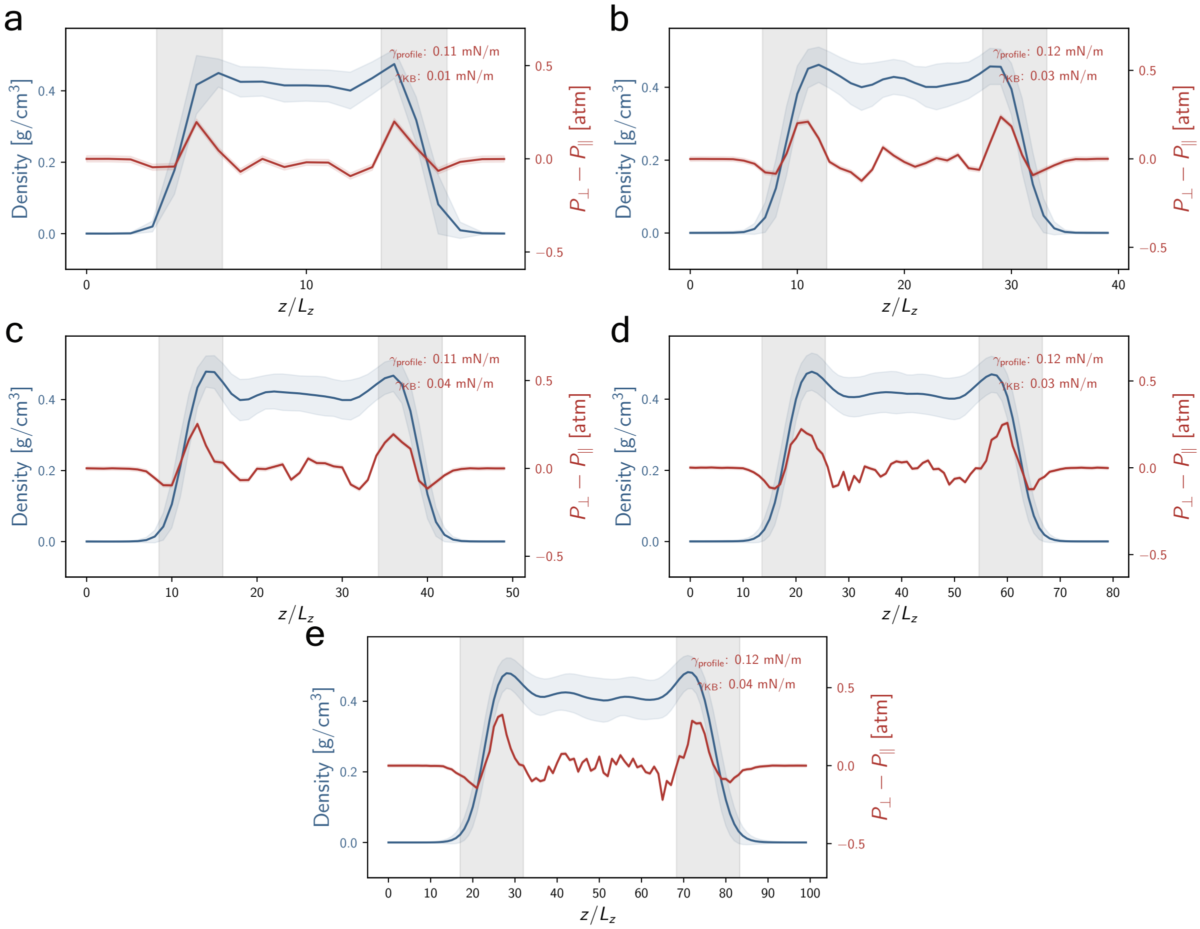


Figure S36. **Stress Profiles for slab simulations of DDX4 condensates at $0.9T_c$ demonstrating invariance to the number of bins used in the calculation of surface tension.** These stress profiles also show the density profile and surface tension value predicted by the Kirkwood–Buff formalism applied in LAMMPS.

A temperature dependence of the form: $\gamma \approx (T_c - T)^{1.26}$ corresponding to the three-dimensional Ising universality class [4] can be fitted to the calculated surface tension values from simulations to assess their prediction of critical temperatures.

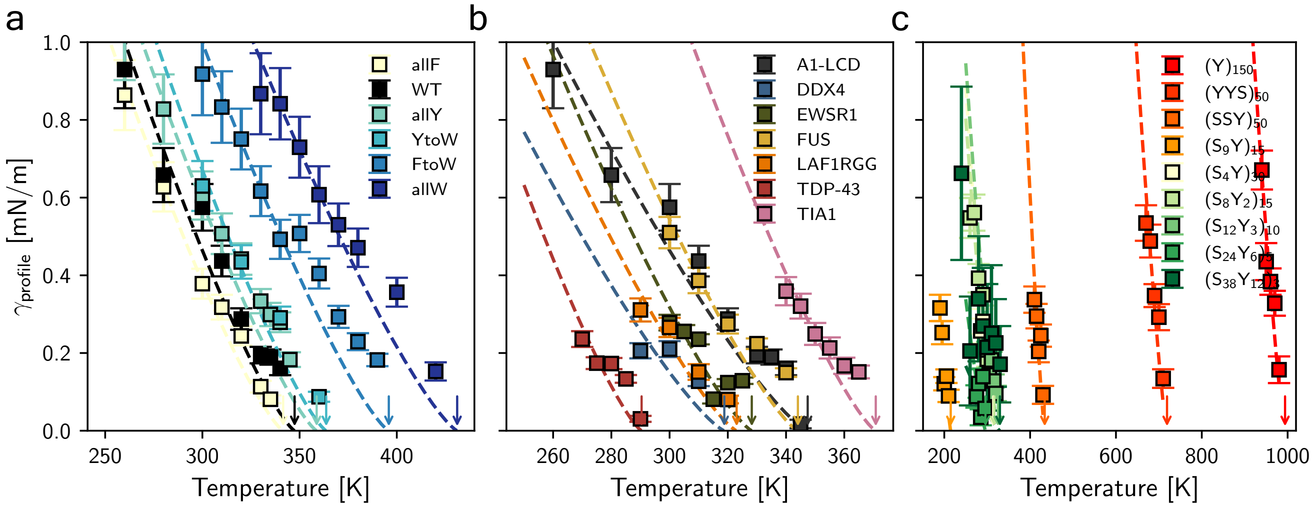


Figure S37. **Temperature dependence of the surface tensions of all studied condensates.** Data is fitted to the analytical form prediction from the 3D Ising universality class.

*Quantifying condensate mobility through molecular diffusion*

To gain insight into how different IDP sequence chemistries and different temperatures affect condensate dynamics we calculate the **diffusion coefficients** of the proteins using the mean squared displacement (MSD) procedure [5] far from the ballistic regime using the Einstein relation.

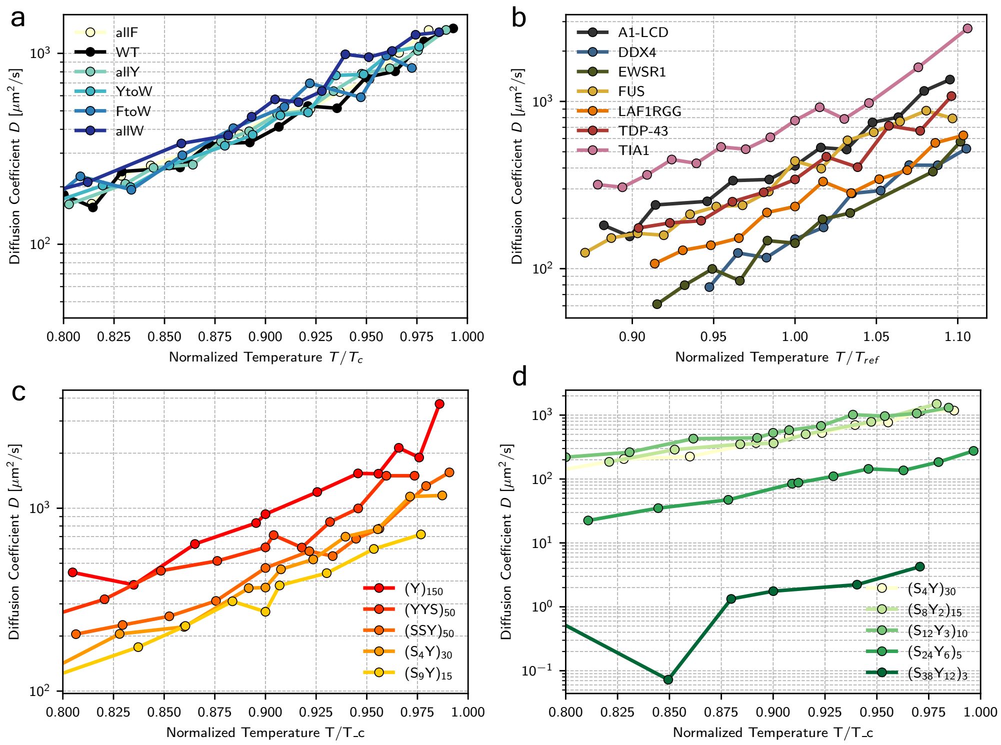


Figure S38. **MSD-derived diffusion coefficients for the studied condensates over normalized temperature.**

We further confirm the correlation between viscosity and diffusion coefficients follows the prediction by the Rouse model [6] for a range of temperatures.

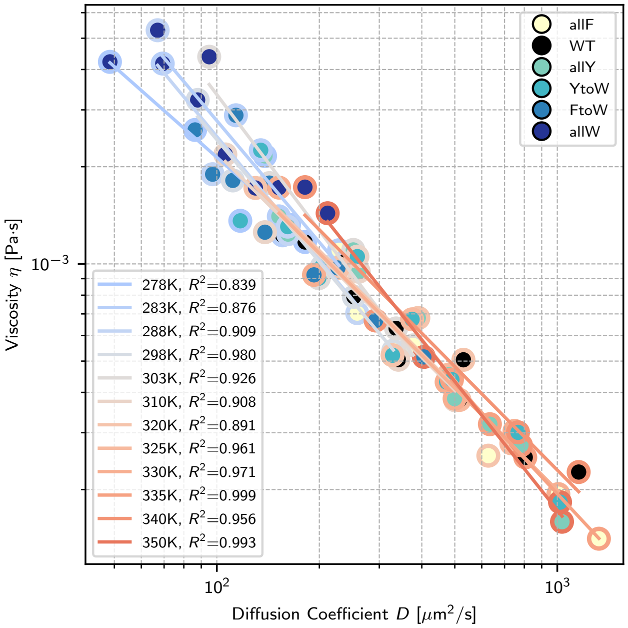


Figure S39. **Relationship between condensate bulk viscosity and bulk diffusion coefficients for A1-LCD variants over a range of temperatures.** The relationship follows the prediction of the Rouse Model.


[1] K. Silmore, M. Howard, and A. Panagiotopoulos, Vapour–liquid phase equilibrium and surface tension of fully flexible lennard–jones chains, Mol Phys (2017).

[2] D. T. Walton, J. Rowlinson, and J. Henderson, The pressure tensor at the planar surface of a liquid, Molecular physics (1983).

[3] A. P. Thompson, H. M. Aktulga, R. Berger, D. S. Bolintineanu, W. M. Brown, P. S. Crozier, P. J. in 't Veld, A. Kohlmeyer, S. G. Moore, T. D. Nguyen, R. Shan, M. J. Stevens, J. Tranchida, C. Trott, and S. J. Plimpton, Lammps - a flexible simulation tool for particle-based materials modeling at the atomic, meso, and continuum scales, Computer Physics Communications **271**, 108171 (2022).

[4] J. M. Prausnitz, R. N. Lichtenthaler, and E. G. D. Azevedo, *Molecular thermodynamics of fluid-phase equilibria* (Pearson Education, 1998).

[5] I.-C. Yeh and G. Hummer, System-size dependence of diffusion coefficients and viscosities from molecular dynamics simulations with periodic boundary conditions, The Journal of Physical Chemistry B **108**, 15873 (2004).

[6] M. Rubinstein and A. N. Semenov, Dynamics of entangled solutions of associating polymers, Macromolecules **34**, 1058 (2001).
